# *Glomage*: A multimodal platform for high-content morphological and RNA profiling of glomeruli in zebrafish and mouse models

**DOI:** 10.1101/2025.10.31.684126

**Authors:** Maximilian Schindler, Tim Lange, Sophia-Marie Bach, Soeren S. Lienkamp, Nicole Endlich

## Abstract

Chronic kidney disease affects over 800 million people worldwide, but therapeutic options remain limited. The glomerulus, with its highly specialized podocytes, forms the functional core of blood filtration. However, studying this complex three-dimensional kidney compartment is difficult, which has slowed progress in identifying drugs. Zebrafish larvae are a powerful model for kidney research and drug discovery, but histological staining of individual larvae is time-consuming, and scalable methods for glomerular analysis have been lacking. Here, we establish a high-throughput workflow for isolating, staining, imaging, as well as extracting RNA from hundreds of zebrafish glomeruli simultaneously. The workflow preserves the three-dimensional architecture, enables quantitative assessment of podocyte numbers, and was validated in a podocyte-specific injury model. Moreover, we adapted the protocol for mouse glomeruli, allowing reliable detection of the age-related loss of podocytes. *Glomage* provides a systematic, cross-species platform for glomerular analysis, opening new opportunities for kidney research and drug discovery.

## Introduction

Chronic kidney disease affects an estimated 10% of the global population and is a leading cause of morbidity and mortality worldwide.^1^ Its prevalence continues to rise due to aging populations, hypertension, and diabetes, highlighting the urgent need to understand how morphological and gene expression changes impact kidney function.^2^ The glomeruli, highly specialized filtration units of the kidney, consist of parietal epithelial cells, mesangial cells, fenestrated capillary endothelia, the glomerular basement membrane, and morphologically complex podocytes that together ensure effective blood filtration. Podocytes represent a unique population of postmitotic epithelial cells, that envelop the glomerular capillaries with interdigitating foot processes. By forming the slit diaphragm and expressing key proteins such as nephrin, together with transcription factors like Dach1, they play a pivotal role in maintaining the size- and charge selectivity of the glomerular filtration barrier.^3–5^ Despite its critical role, the glomerular structure remains difficult to study in depth. Traditional histological approaches using paraffin- or cryosections are indispensable for assessing morphological changes and protein expression, yet they provide only two-dimensional snapshots of a three-dimensional structure. Moreover, loss of podocytes, whether through apoptosis or detachment, is irreversible and is a major driver of disease progression, making podocyte number a crucial indicator of kidney health and a strong predictor of renal outcomes. However, conventional methods may under- or overestimate podocyte counts due to inherent sampling bias.^6–8^ In contrast to classical methods, advanced imaging approaches such as optical clearing combined with confocal or light-sheet microscopy have enabled whole-glomerulus visualization but are time-intensive, technically demanding, and require specialized equipment, limiting their scalability.^9–11^ These challenges have slowed progress towards standardized, high-throughput and deep glomerular analysis.

Zebrafish larvae have emerged as a versatile vertebrate model for kidney biology and disease due to their rapid development, optical transparency, and genetic accessibility.^12–14^ Furthermore, approximately 70% of human genes have at least one zebrafish ortholog, emphasizing its utility as a model for human disease.^15^ The larval zebrafish pronephros closely mimics the architecture and filtration function of the mammalian glomerulus, making it a powerful model for studying e.g. podocytopathies and to identify drugs to treat kidney disease.^16–22^

The zebrafish pronephros consists of only a single glomerulus attached to a pair of tubules. While tissue sections and whole-mount imaging of single zebrafish larvae provide valuable insights, they are not yet suitable for high-throughput applications.^23^ Molecular studies face similar challenges: isolating sufficient, pure glomerular RNA from zebrafish larvae has been technically difficult and time-consuming, so most transcriptomic analyses rely on whole-larva samples, masking glomerular or podocyte-specific expression patterns.^24,25^

To address these limitations, we developed a protocol for batch isolation of intact and whole larval zebrafish glomeruli, enabling fast and scalable immunofluorescence, 3D-high-resolution imaging, an unbiased and automated podocyte count as well as downstream molecular analyses. This protocol was applied in a larval model of focal segmental glomerulosclerosis (FSGS), where it effectively identified structural damage, enabled a reliant quantification of podocyte loss and detection of volumetric changes. The same staining and quantification protocol was successfully applied to isolated mouse glomeruli, confirming the age-dependent podocyte loss.

In summary, this protocol enables scalable isolation and analysis of whole glomeruli across animal and disease models, supporting advances in glomerular research, biomarker discovery, and drug screening.

## Results

### *Glomage*: High-throughput isolation of larval zebrafish glomeruli

To enable high-throughput analysis of pronephric zebrafish glomeruli, we developed a protocol for their batch isolation and immunofluorescent labeling. For this purpose, we generated a *nphs2*:eGFP (*Podo:GFP*) zebrafish line, which expresses enhanced green fluorescence protein (eGFP) endogenously and exclusively in glomerular podocytes (Fig. S1). The first step of the protocol is the fixation of the larvae at 5 days post fertilization (dpf) in 2% paraformaldehyde (PFA) for 90 min at room temperature (RT), which is a critical step preceding tissue dissociation. The procedure was carried out using 100 fixed eGFP-positive larvae as the standard sample size. The larvae were placed in a tube together with ∼100 µl ceramic beads with a size of 1 mm and were dissociated with a Fast-Prep-24^TM^ device for 13 seconds with 4 m/s, resulting in the release of glomeruli (Fig. 1A). The efficiency of dissociation is highly dependent on the fixation parameters such as PFA concentration, incubation time and temperature as well as dissociation intensity and time. Insufficient fixation or excessive dissociation lead to glomerular damage, compromising tissue integrity. Moreover, extended fixation times can result in reduced glomerular yield and in a loss of antigenicity of native tissue as it was also described in the literature.^26,27^ To prevent adhesion of isolated glomeruli to plastic surfaces, all materials were precoated with fetal bovine serum (FBS) since glomeruli are highly charged.^28,29^ Following larval dissociation, intact glomeruli were collected under a fluorescence stereomicroscope using a gel-loading pipette and subsequently transferred onto a cell culture insert with a polyethylene terephthalate (PET) membrane (mesh size: 8 µm, Greiner ThinCert®). The inserts were transferred into a 24-well plate, in which the immunofluorescence staining was performed. The staining procedure was based on previously established protocols with minor modifications,^16,30,31^ which are described in detail in the methods section. The minimum volume of each (antibody) solution was 500 µl per well to ensure that the PET membrane carrying glomeruli remained fully covered. At the end of the staining procedure, the insert membrane was excised with a scalpel and mounted onto a glass slide for microscopy (Fig. 1A). For automated imaging, low-magnification overview maps (1.25x objective) were acquired to facilitate navigation to individual glomeruli with higher-magnification objectives (20x or 60x, Fig. 1B-D).

**Figure 1:**
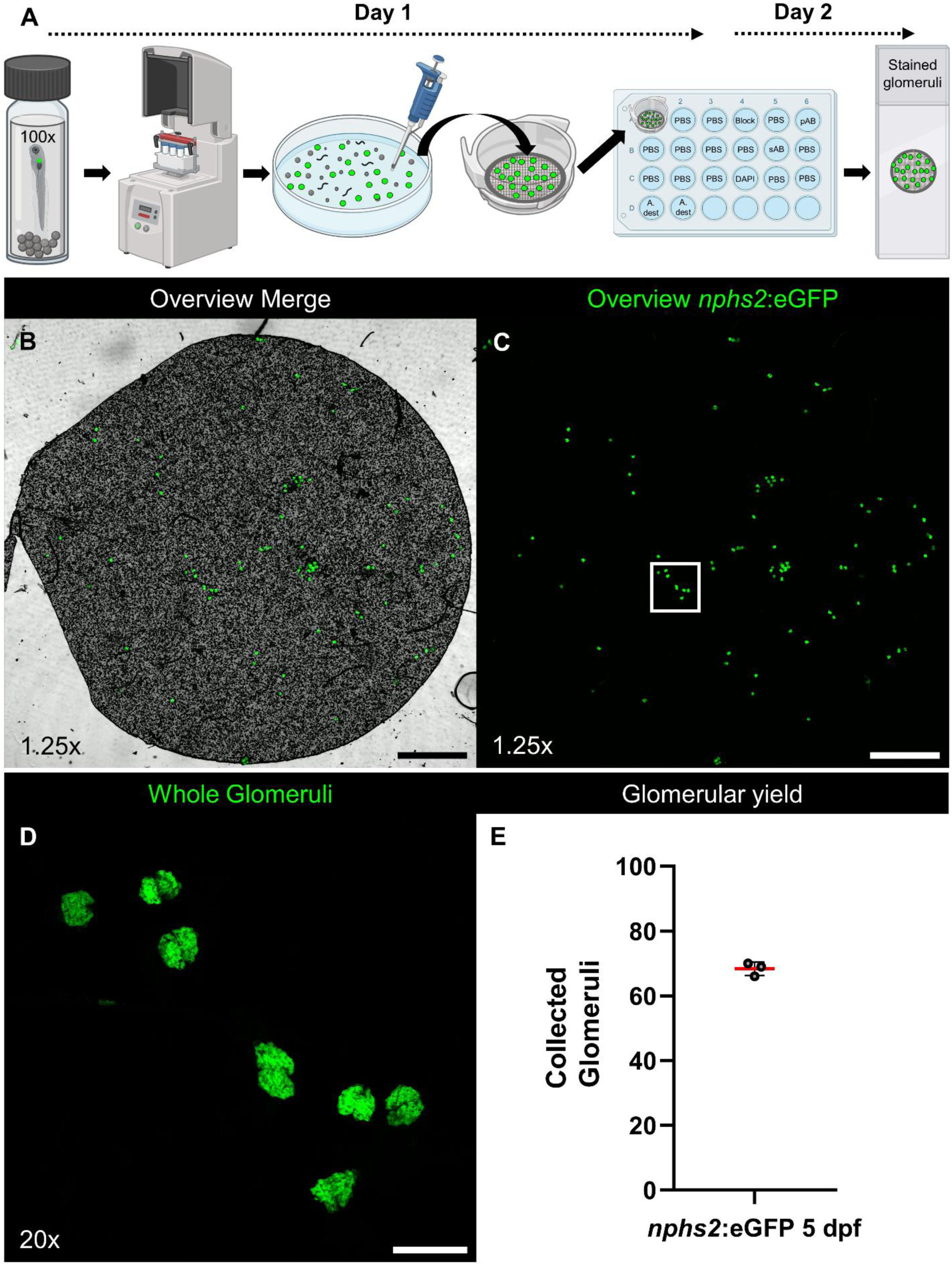
Batch isolation of whole zebrafish glomeruli. PFA-fixed *Podo:GFP* larvae are dissociated using ceramic beads, and fluorescent, intact glomeruli are collected and transferred to mesh baskets. Immunofluorescence staining is performed in a 24-well plate, and the cut mesh containing the stained glomeruli is mounted onto a glass slide (**A**). An overview image acquired with a 1.25x objective shows the mesh in brightfield and the fluorescent glomeruli (**B**). The corresponding eGFP-only image illustrates all glomeruli (**C**). A higher magnification using a 20x objective reveals intact, peanut-shaped larval zebrafish glomeruli (**D**). Using 100 larvae per isolation resulted in a mean of 68 (SD: 2, n=3) isolated, stained, and mounted whole zebrafish glomeruli per round (**E**). Scale bars: 1 mm (**B**, **C**), 100 µm (**D**).

At the end of the protocol, 3 sets of 100 larvae were used as input. Manual counting of glomeruli on PET membranes resulted in 68 (SD: 2) isolated, stained, and mounted whole glomeruli. Given that zebrafish larvae only possess a single glomerulus, this corresponds to a 68% recovery rate.

### *Glomage*: A platform for 3D immunofluorescence analysis of larval glomeruli

For antibody staining, we consistently used antibodies and working concentrations that had already been validated and optimized in established protocols for zebrafish larval cryosections (Fig. S2). They were therefore expected to perform similarly in native, non–paraffin-embedded whole glomeruli.

To demonstrate the applicability of this protocol, we used antibodies against podocyte-specific proteins, such as the slit diaphragm proteins nephrin and podocin; the transcription factor Pax2a, which is specific for parietal epithelial cells (PECs) and proximal tubule epithelial cells (PTECs); Ehd3, a marker of glomerular endothelial cells; and laminin, a major component of the glomerular basement membrane (GBM). Positive antibody signals alongside endogenous eGFP expression and nuclei can be seen in single optical sections of the stained glomeruli (Fig. 2A-E, Movie S1).

**Figure 2:**
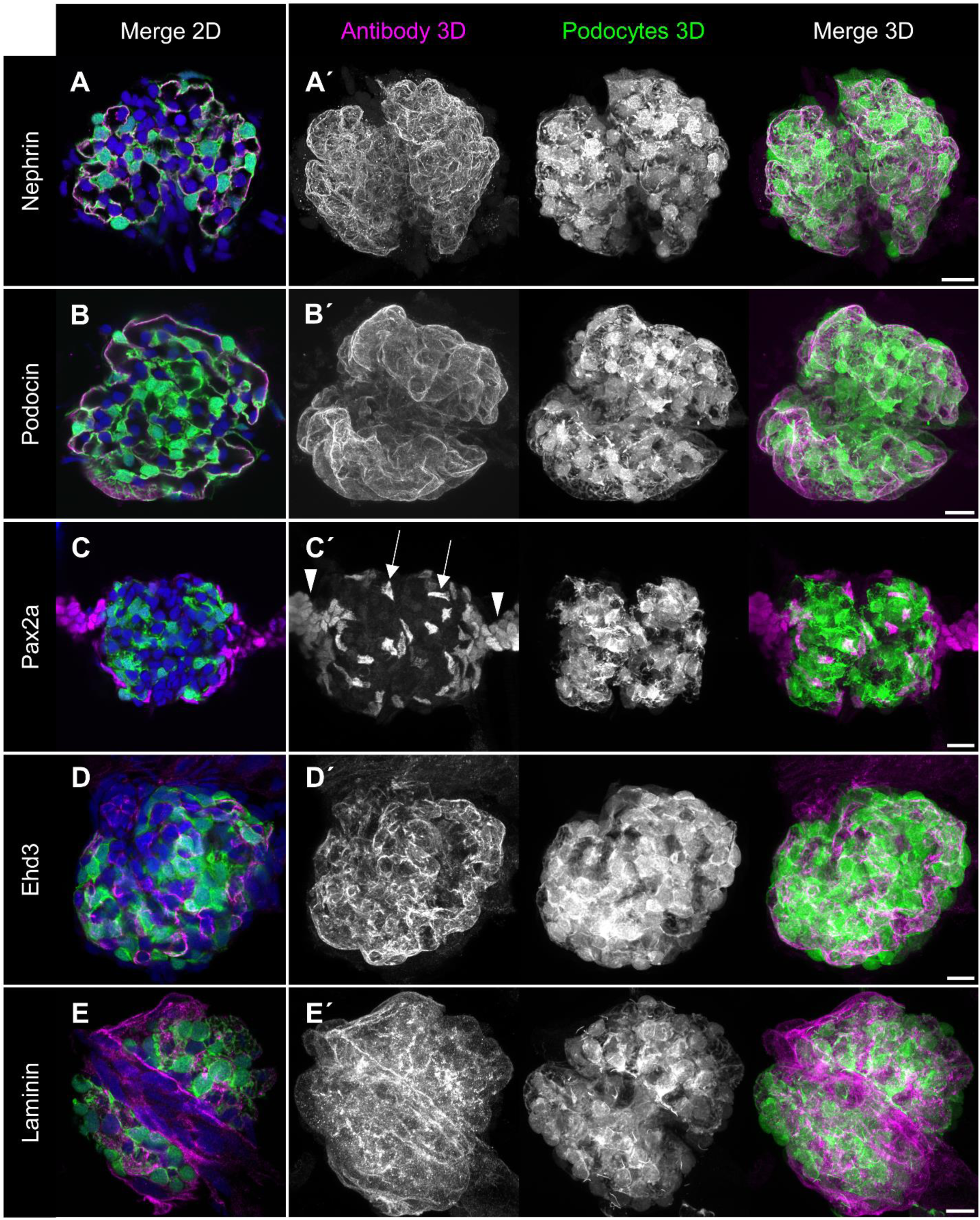
Visualization of whole zebrafish glomeruli using different antibodies in the *Podo:GFP* strain. Antibodies against the slit diaphragm proteins nephrin and podocin show a continuous, linear staining along the filtration slits in single optical sections (**A**, **B**) and in corresponding maximum intensity projections (3D, **A′**, **B′**). Staining with a Pax2a antibody reveals the coverage of the glomerulus by nuclei of parietal epithelial cells (arrows), and identifies two populations of proximal tubule cells connected to the glomerulus (arrowheads, **C**, **C′**). An Ehd3 antibody specifically labels fenestrated endothelial cells (**D**, **D′**), while the laminin staining highlights the basement membranes (**E**, **E′**). Podocytes are endogenously labeled with eGFP (green), nuclei (blue) are displayed in the 2D panels and omitted in the 3D merge panels. Scale bars: 10 µm.

For each glomerulus, optical z-stacks were acquired at 1 µm intervals, yielding a volumetric range of 25–60 µm, followed by maximum intensity projections (MIPs). This approach enabled 3D reconstruction of the whole slit diaphragm architecture in larval zebrafish glomeruli (Figure 2A′, B′, Movie S2 and S3). This procedure also revealed the localization of Pax2a-positive PECs surrounding the glomerulus. Pax2a also labelled PTECs of the two attached proximal tubules, which in some cases remained connected to the glomeruli (Fig. 2C′). In addition, the staining visualized Ehd3-positive, fenestrated endothelial cells within the glomeruli consistent with their known expression pattern (Fig. 2D′).^32^ Additionally, labeling of F-actin with fluorescence-conjugated phalloidin showed the brush border of the convoluted proximal tubule in 3D, including the short neck segment which connects the glomerulus to the tubule. The neck segment was characterized by reduced apical microvillus density (Fig. S3, Movie S4).

Taken together, all five antibodies produced specific and reproducible staining patterns in whole-mount glomeruli, demonstrating that this protocol enables, for the first time, rapid and efficient immunolabeling of multiple glomerular target structures across large numbers of intact whole glomeruli.

### *Glomage*: Rapid quantification of total podocyte number in whole larval glomeruli

Because podocyte number changes with both disease progression and organismal age, precise quantification of these postmitotic cells is essential. A reduction below a critical threshold compromises glomerular and renal function, ultimately resulting in proteinuria, the leakage of high-molecular-weight proteins into the urine.^33^ To establish this analysis, z-stacks of isolated glomeruli from *Podo:GFP* larvae were analyzed. Since eGFP is a protein of less than 60 kDa and can therefore diffuse into the nucleus, podocyte nuclei can be readily identified by their exclusive nuclear labelling with eGFP (Fig. 3A-D). To address the challenge of three-dimensional nuclear segmentation, particularly the risk of counting the same nucleus multiple times across optical slices, an AI-based image analysis software was used to segment and quantify eGFP-positive nuclei. By applying an eGFP intensity threshold after segmentation of all nuclei within and surrounding each glomerulus, it was possible to reliably determine the number of podocytes per glomerulus from full z-stacks. Analysis of 18 glomeruli revealed an average of 88.40 podocytes per glomerulus (SD: 13.23) in zebrafish larvae at 5 dpf (Fig. 3E-H).

**Figure 3:**
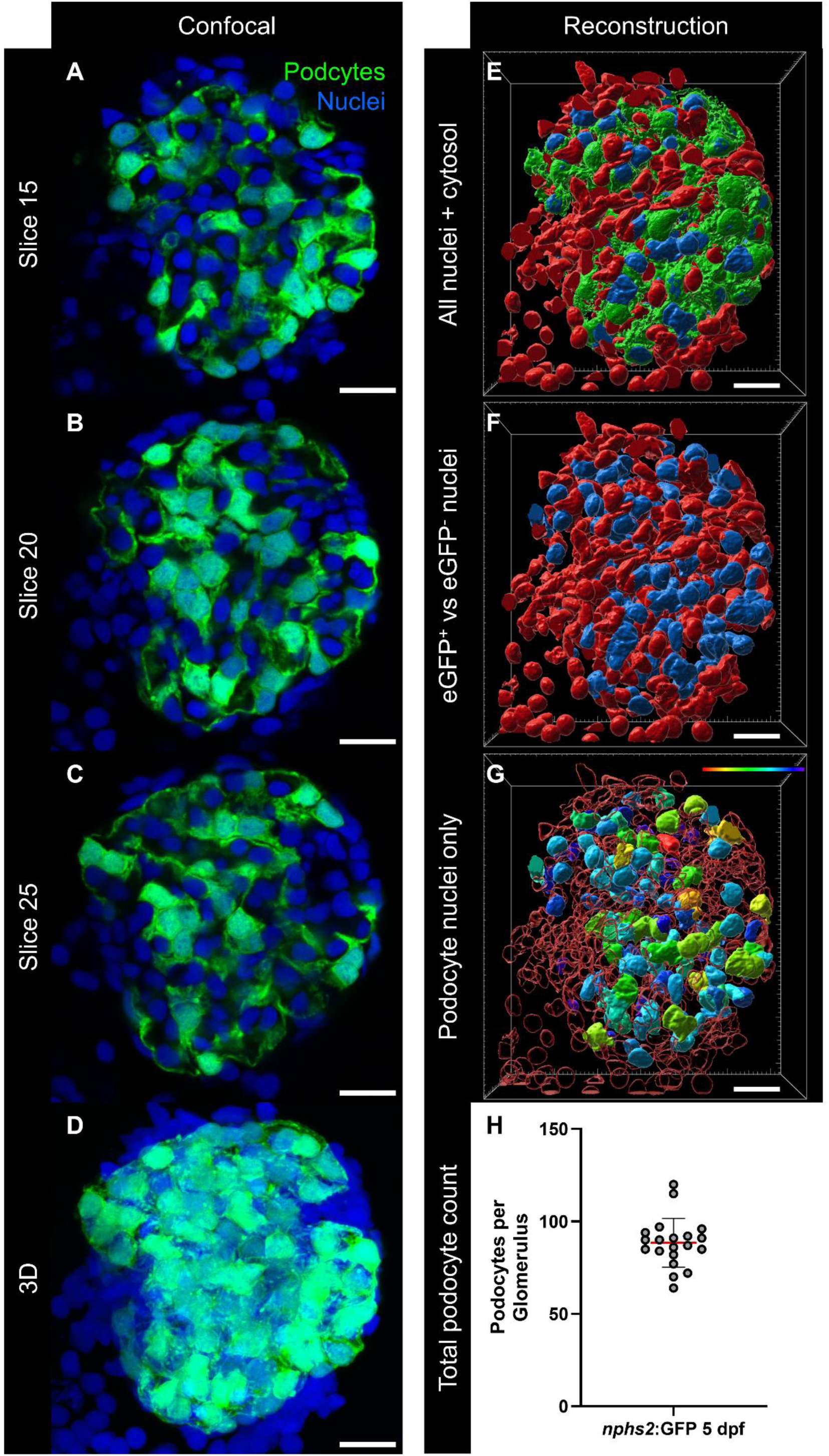
Establishment of the total podocyte count. Confocal image stacks of isolated zebrafish glomeruli show eGFP-positive podocyte nuclei in individual optical sections (**A–C**) and in the corresponding projection (**D**). AI-based segmentation visualizes the podocyte cytoplasm (green), eGFP-positive nuclei (blue), and eGFP-negative nuclei (red) (**E**). Panel **F** shows the spatial distribution of eGFP-positive (blue) and eGFP-negative (red) nuclei within the glomerulus. All eGFP-positive nuclei are used to define the total number of podocytes in a single glomerulus (**G**). Podocyte nuclear volume is color-coded, ranging from 6.7 µm³ (purple/blue) to 83.3 µm³ (red). Automated analysis of n=18 zebrafish glomeruli at 5 days post fertilization (dpf) revealed an average of 88.40 podocytes per glomerulus (SD: 13.23) (**H**). Scale bars: 10 µm.

Therefore, *Glomage* allows for the first time the robust and scalable quantification of podocyte number in transgenic zebrafish larvae without the use of antibodies. Accidental co-isolation of adjacent tissue does not interfere with the analysis, as only podocytes express eGFP in this specific zebrafish strain.

### *Glomage*: Podocyte quantification in podocyte injury models

To evaluate the functional applicability of this novel podocyte quantification method, we used the *nphs2*:NTR-mCherry (*Cherry*) transgenic zebrafish strain, which expresses the bacterial enzyme nitroreductase (NTR) and mCherry exclusively in podocytes under the control of the *podocin* promoter (Fig. S1).^34^ Upon exposure of zebrafish larvae to 50 nM nifurpirinol (NFP) in the tank water at 4 dpf, podocyte injury develops as NFP is converted into a toxin by the NTR within podocytes, mimicking key features of mammalian FSGS.^30,31,35,36^ Since mCherry fluorescence in podocytes was markedly reduced after 24 hours of NFP treatment, experiments were terminated at this time point to avoid complete podocyte loss.^16^ We analyzed three groups: (i) a control group treated with 0.2% dimethyl sulfoxide (DMSO), (ii) an injury group treated with 50 nM NFP, and (iii) a group co-treated with NFP and the small molecule NM_187 to prevent disease progression. After 24 hours, the *Glomage* procedure was performed, and glomeruli were stained for the slit diaphragm protein podocin. 3D-image analysis of healthy control glomeruli displayed a continuous, meandering pattern of podocin along the glomerular filtration barrier and a homogeneous mCherry fluorescence in podocytes (Fig. 4A). In contrast, NFP treatment induced clear morphological signs of podocyte injury, including a punctate slit diaphragm staining pattern, pseudocyst formation, intracellular vacuoles, and a fragmented or inhomogeneous mCherry signal (Fig. 4B). Co-treatment with NM_187 fully rescued these features: Podocin staining returned to a linear pattern, and mCherry fluorescence was homogeneously distributed (Fig. 4C). Our automated quantification pipeline was successfully adapted to work with mCherry instead of eGFP as a nuclear marker, confirming its fluorophore-independence. Analysis of podocyte nuclear volume revealed that NFP treatment significantly reduced the average nuclear volume (mean: 33.44 µm³, SD: 2.4) compared to controls (mean: 48.46 µm³, SD: 2.82). Co-treatment of the larvae with NM_187 significantly restored nuclear volume (mean: 39.81, SD: 1.84, Fig. 4D). A deeper analysis of individual optical slices revealed abundant small mCherry-positive nuclear fragments in the NFP group, representing apoptotic bodies. These were absent in the control and NM_187 co-treatment groups (Fig. S4). To exclude apoptotic fragments from the unbiased podocyte count, a volume threshold was set at one standard deviation below the mean nuclear volume of the control group. The same threshold was applied consistently across all groups and all nuclei exceeding this value were defined as nuclei from viable podocytes. After applying the threshold, podocyte counts in healthy *Cherry* larvae (mean: 95.6; SD: 12.90) closely matched those in *Podo:GFP* larvae (mean: 88.40, SD: 13.23, Fig. 3H). NFP treatment resulted in a significant loss of viable podocytes (mean: 48.2, SD: 12.14), whereas co-treatment with NM_187 fully prevented podocyte loss, with final counts comparable to healthy controls (mean: 95.30, SD: 19.72, Fig. 4E).

**Figure 4:**
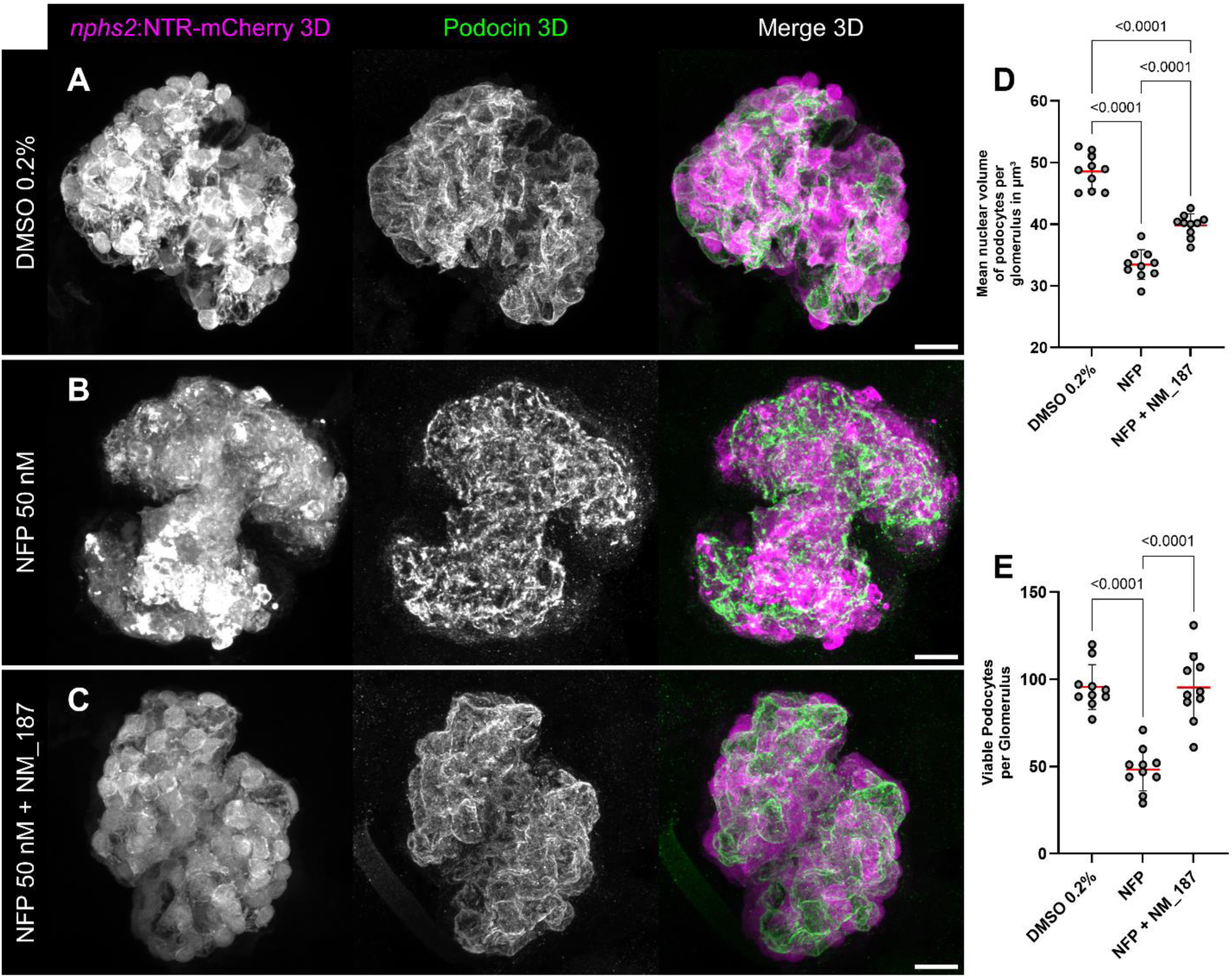
Histology and podocyte counts in healthy, NFP-treated, and NFP + NM_187-treated *Cherry* larvae. Healthy control larvae expressing mCherry and the nitroreductase (NTR) in podocytes show a linear Podocin staining pattern along the slit diaphragm (**A**). Targeted podocyte injury with NFP for 24 hours induces structural damage and disruption of the slit diaphragm (**B**). Treatment with the compound NM_187 prevents podocyte injury and preserves both podocyte integrity and the linear Podocin expression pattern (**C**). Podocyte injury significantly reduces nuclear volume, which is rescued by NM_187 treatment (DMSO mean: 48.46 µm³, SD: 2.82, NFP mean: 33.44 µm³, SD: 2.4), NFP + NM_187 mean: 39.81, SD: 1.84, D). Automated AI-based segmentation and quantification reveal a significant reduction in the number of viable podocytes per glomerulus following specific injury (mean: 48.2; SD: 12.14), compared to healthy DMSO-treated controls (mean: 95.6; SD: 12.9). Treatment with NM_187 completely prevents podocyte loss (mean: 95.3; SD: 19.72) (E). Statistical analysis was performed with n=10 glomeruli in each group using one-way ANOVA followed by Tukey’s multiple comparisons test and p≤0.05 was considered statistically significant. Scale bars: 10 µm.

This is the first demonstration that differences in podocyte number in zebrafish larvae can be determined in injury and in response to treatment with a drug that inhibits the development of larval FSGS, highlighting the potential of this approach for drug screening experiments and toxicity assessments.

### *Glomage*: Volumetric analysis of healthy and injured larval glomeruli

To further assess the morphological impact of NFP and the potential protective effect of NM_187 on podocyte structure, we performed volumetric analysis of glomerular podocyte cytoplasm in whole glomeruli of *Cherry* larvae. Confocal z-stacks were used to render three-dimensional, contiguous volumes engulfed by mCherry-positive podocyte structures (Fig. 5A-C’). Quantitative analysis revealed a significant reduction in glomerular mCherry volume in NFP-treated larvae (mean: 26,41 µm³, SD: 4,89) compared to controls (mean: 38,05 µm³, SD: 4,00). Notably, co-treatment with NM_187 significantly restored podocyte volume to near-control levels (mean: 35,58 µm³, SD: 8,07; Fig. 5D). In addition to volume changes, the sphericity of glomeruli was assessed in which a value of 1 describes a perfect sphere. The sphericity of the mCherry-positive volume was significantly reduced in both NFP (mean: 0.44, SD: 0.05) and NFP + NM_187-treated larvae (mean: 0.51, SD: 0.07) relative to controls (mean: 0.60, SD: 0.08; Fig. 5E), indicating a loss of normal glomerular morphology that was only partially rescued by NM_187.

**Figure 5:**
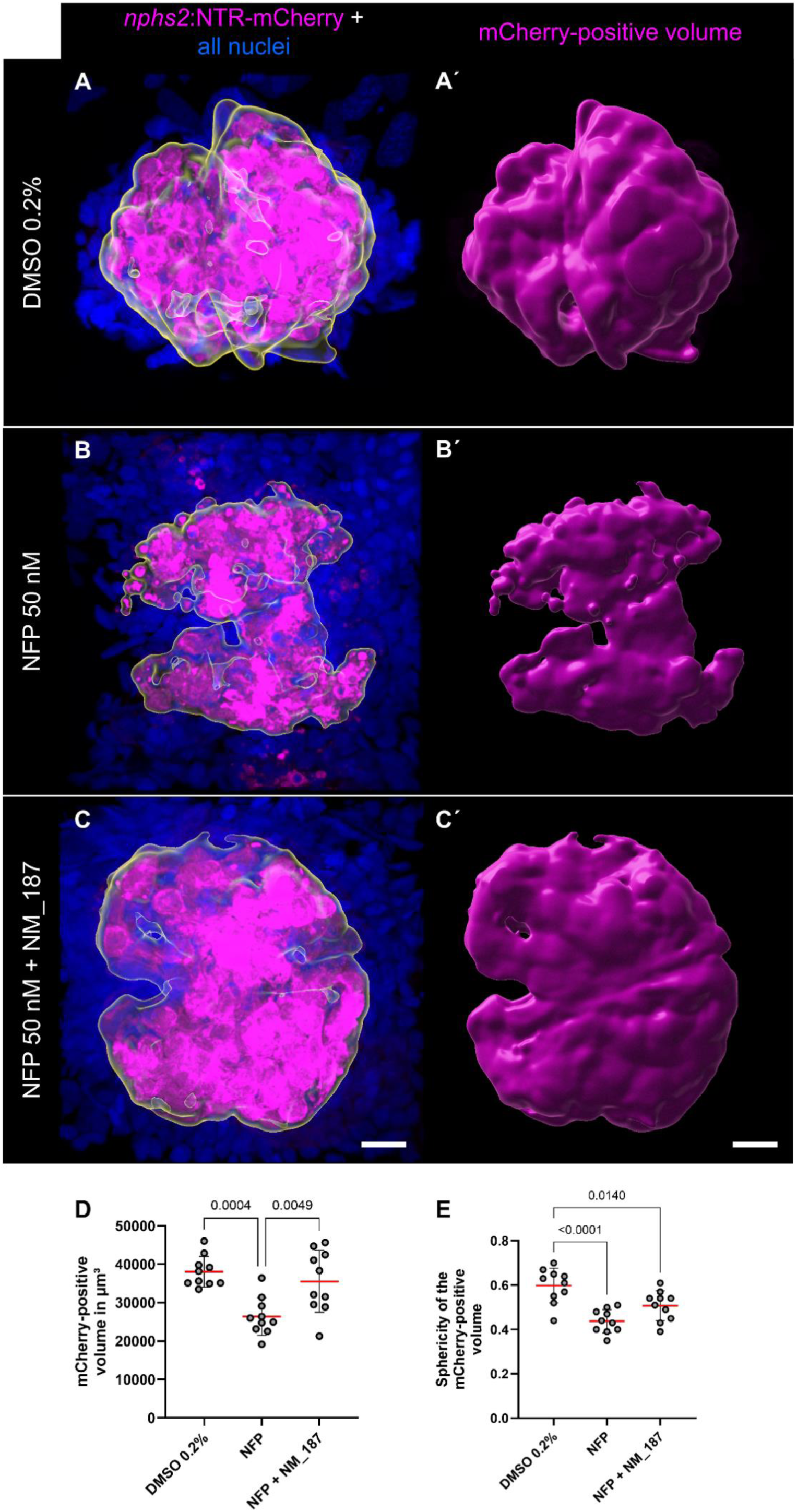
Volumetric depiction and analysis of DMSO, NFP and NFP + NM_187-treated *Cherry* larvae. The continuous mCherry-positive podocyte cytoplasm in glomeruli from confocal z-stacks (**A**–**C**) was used to render a three-dimensional, contiguous volume (**A′**–**C′**). Quantification revealed a significant reduction in mCherry-positive volume in NFP-treated larvae (mean: 26,406 µm³, SD: 4,889) compared to DMSO controls (mean: 38,053 µm³, SD: 4,004). Co-treatment with NM_187 significantly restored mCherry volume to near-control levels (mean: 35,580 µm³, SD: 8,070; **D**). Glomerular sphericity analysis showed a significant reduction in roundness in both NFP (mean: 0.44, SD: 0.05) and NFP + NM_187-treated larvae (mean: 0.51, SD: 0.07) compared to controls (mean: 0.60, SD: 0.08, **E**). For each group, n=10 whole glomeruli were analyzed. Statistical analysis was performed using one-way ANOVA followed by Tukey’s multiple comparisons test. p≤0.05 was considered statistically significant. Scale bars: 10 µm.

### *Glomage*: Isolation of pure glomerular RNA

In zebrafish larvae, isolating single organs in large numbers within a short time frame and with sufficient purity remains challenging. As a result, most studies rely on whole-larva analysis. To expand the utility of our batch glomerulus isolation protocol beyond imaging applications, we developed a modified workflow to obtain pure glomerular mRNA. In this approach, larvae from the *Cherry* strain were dissociated without PFA fixation under mild conditions (1 s with 4 m/s), and glomeruli were manually selected under fluorescence guidance. Several wash steps and stringent visual quality control were applied to eliminate adjacent or contaminating cells, which are critical for RNA purity. RNA was extracted from 50 or 100 isolated glomeruli using a single-cell RNA isolation kit, followed by complementary DNA (cDNA) synthesis and RT-PCR as well as RT-qPCR analysis (Fig. 6A). Gene expression was assessed using primers for *eef1a1a* (housekeeping gene), *nphs2* (podocyte-specific), and *mCherry* (transgene, podocyte-specific). To assess sample purity, we included markers specific to other organs: *gnat1* (eye-specific), *myl7* (heart-specific), and *ctrb1* (pancreas-specific).

**Figure 6:**
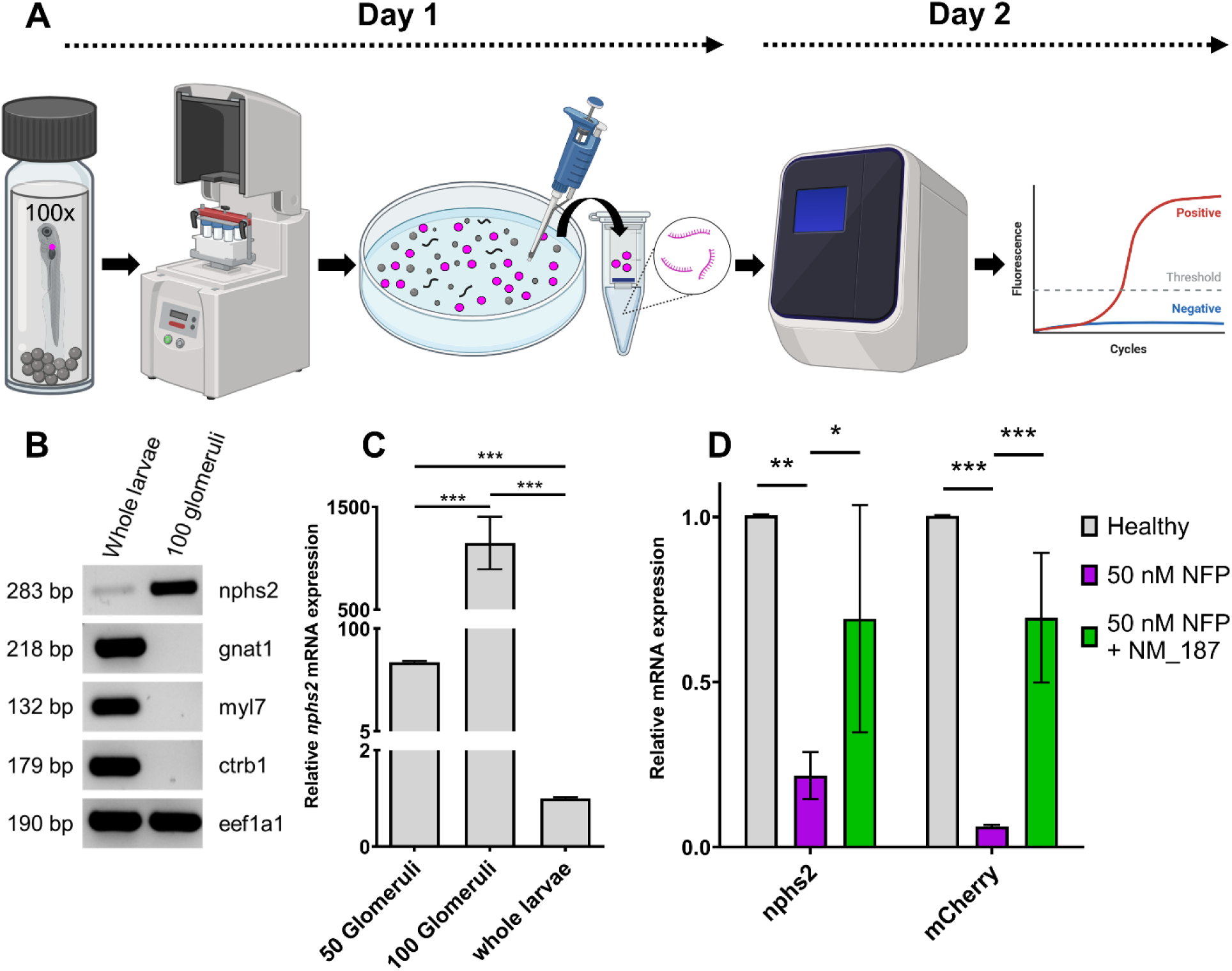
Quantitative glomerular RNA isolation and downstream application in podocyte injury. Unfixed *Cherry* larvae were dissociated and glomerular RNA was isolated using a single-cell RNA kit at 5 dpf. Subsequent cDNA synthesis and RT-PCR as well as RT-qPCR was performed using podocyte and non-podocyte specific primers (**A**). Primers detecting *nphs2* (*podocin*, podocyte-specific), *mCherry* (podocyte-specific) and *gnat1* (eye), *myl7* (heart) *ctrb1* (pancreas) were used to assess sample purity. cDNA from whole larvae were compared to cDNA derived from 100 isolated glomeruli. RT-PCR results show absence of non-glomerular markers (*gnat1*, *myl7*, *ctrb1*) in isolated glomeruli, while *nphs2* is strongly enriched compared whole-larva levels (**B**). RT-qPCR shows that *nphs2* levels are substantially higher in isolated glomeruli, showing a 68.9-fold change (SD: 1.1) in 50 glomeruli and 1148.8-fold (SD: 256.7) in 100 glomeruli compared to whole larvae (1.0-fold, SD: 0.03, **C**). In NFP-induced injury, *nphs2* (0.22-fold, SD: 0.06) and *mCherry* (0.06-fold,SD: 0.00) are downregulated. Co-treatment with NM_187 significantly restores expression (*nphs2*: 0.69-fold, SD: 0.29, *mCherry*: 0.70-fold, SD: 0.19, **D**). For RT-qPCR experiments, three independent isolations were compared (n=3). Statistical analysis was conducted by using two-way ANOVA followed by Bonferroni-post-hoc test. p≤0.05 was considered statistically significant.

In conventional RT-PCR (Fig. 6B, Fig. S5), the non-glomerular markers *gnat1*, *myl7*, and *ctrb1* were not detectable in isolated glomerular RNA, whereas they were readily amplified from whole-larva samples. This confirms that the isolation workflow yields highly pure glomerular material without detectable contamination from other organs. In contrast, the podocyte-specific marker *nphs2* was robustly expressed in glomerular samples and only slightly expressed in whole larvae samples, validating the specificity of the isolation procedure (Fig. 6B, Fig. S5).

RT-qPCR analysis demonstrated a strong enrichment of *nphs2* transcripts in isolated glomeruli compared to whole larvae. The mean relative *nphs2* expression in 50 isolated glomeruli was 68.9-fold (SD: 1.1), and in 100 glomeruli 1148.8-fold (SD: 256.7), whereas whole-larva samples served as baseline (1.00-fold, SD: 0.03, Fig. 6C). These results indicate a more than thousand-fold enrichment of podocyte-specific transcripts in isolated glomeruli while maintaining high RNA purity and absence of non-glomerular gene expression.

To further validate the approach in a functional context, *nphs2* and *mCherry* expression were analyzed in a podocyte injury model. Larvae were treated with 0.2% DMSO, 50 nM NFP, or 50 nM NFP and NM_187. Upon NFP treatment, *nphs2* levels dropped markedly to 0.22-fold (SD: 0.06), and *mCherry* decreased to 0.06-fold (SD: 0.00), consistent with severe podocyte injury and loss of podocyte-specific gene expression compared to the DMSO control. Co-treatment with NFP and NM_187 significantly restored expression of *nphs2* to 0.69-fold (SD: 0.29) and *mCherry* to 0.70-fold (SD: 0.19), indicating a protective or restorative effect of NM_187 on podocyte integrity and transcriptional activity (Fig. 6D).

Together, these results demonstrate that our workflow additionally enables high-purity RNA extraction suitable for sensitive transcriptional analyses and that it can reliably detect transcriptional responses in podocyte injury models and treatments.

### *Glomage*: Application of batch staining to isolated mouse glomeruli

After establishing the staining, imaging, and quantification pipeline in zebrafish larvae, we next adapted the method to isolated mammalian glomeruli. To this end, glomeruli were isolated from *nephrin*:CFP mice, expressing cyan fluorescence protein (CFP) exclusively in podocytes,^37^ by using a sieving and differential adhesion approach which is already published.^38^ Shortly after isolation, glomeruli were fixed in PFA, and their density in suspension was assessed under a stereomicroscope. To avoid overloading the culture inserts and especially the PET membrane, a volume corresponding to approximately 50–100 glomeruli was transferred into each insert. The staining and imaging protocol developed for zebrafish glomeruli was then applied to mouse glomeruli (Fig. 7 A). Low-magnification images of the cut PET membrane revealed a high density (>50) of intact, CFP-positive glomeruli after staining (Fig. 7B, C). Higher magnification MIPs demonstrated successful immunolabeling with an antibody against the slit diaphragm protein nephrin, while CFP fluorescence in podocyte cytosol was well preserved (Fig. 7D, E). Single optical slices from the z-stacks further confirmed tissue integrity, with podocyte foot processes clearly identifiable in the CFP channel and the nephrin signal localized between them (Fig. 7F-I).

**Figure 7:**
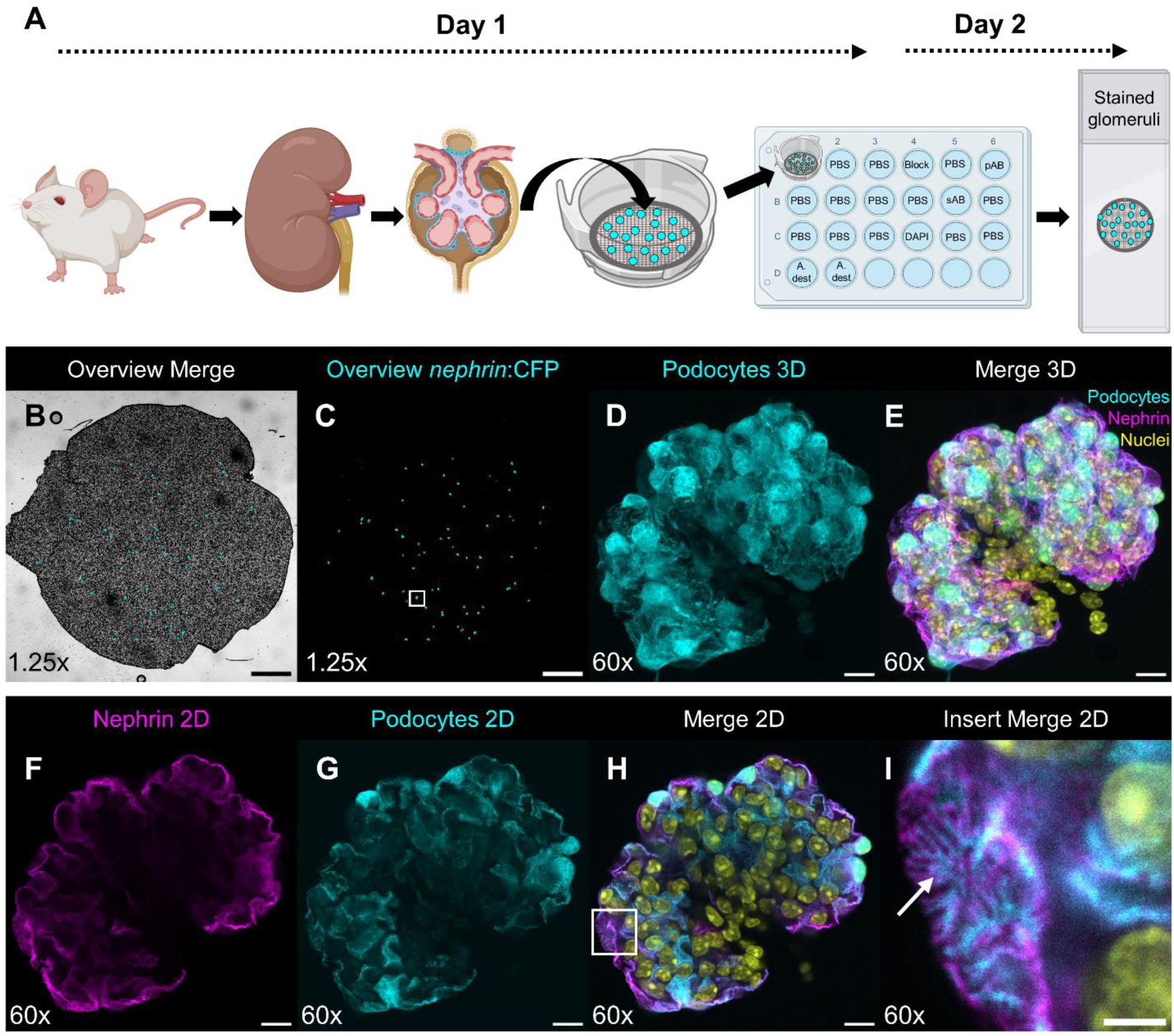
Isolation and staining of whole mouse glomeruli. Glomeruli from n*ephrin*:CFP reporter mice were isolated using a sieve-based protocol combined with differential adhesion, followed by fixation. Fixed glomeruli were transferred to mesh baskets and subjected to immunofluorescence staining in a 24-well plate. The stained mesh was cut and mounted onto a glass slide (**A**). A confocal overview acquired with a 1.25x objective shows the mesh (**B**) with CFP-positive glomeruli (cyan, **C**). Projections of stacks acquired with a 60x objective reveal podocyte cytoplasm (cyan, **D**) and immunostaining for the slit diaphragm protein Nephrin (magenta). Nuclei are shown in yellow (**E**). A single optical section demonstrates the linear Nephrin staining pattern (magenta, **F**) and podocytes in cyan (**G**, **H**). Higher magnification highlights the preserved ultrastructure of the tissue, including resolved podocyte foot processes by confocal microscopy (**I**). Scale bars: 1 mm (**B**, **C**), 10 µm (**D**–**H**), 3 µm (**I**).

Together, these findings demonstrate that the *Glomage* batch-staining and imaging pipeline can be directly transferred to mammalian glomeruli.

### *Glomage*: Total podocyte count in glomeruli of young and aged mice

As a proof of concept, we applied our total podocyte quantification pipeline to isolated, stained glomeruli from 6-month-old (young) and 18-month-old (aged) mice. Podocyte nuclei were identified using an antibody against the transcription factor Dach1, which is highly expressed in adult podocytes, while the slit diaphragm was labeled with an antibody against nephrin in z-stacks of whole glomeruli (Fig. 8A). MIPs of glomeruli from young mice showed continuous nephrin coverage of the glomerular filtration barrier and a high number of Dach1-positive podocyte nuclei (Fig. 8B). In contrast, aged glomeruli exhibited two distinct pathological phenotypes: (1) a mild phenotype characterized by partial nephrin coverage with a punctate staining pattern and a reduced number of Dach1-positive podocytes (Fig. 8C); (2) a severe phenotype with a nearly complete loss of the nephrin signal as well as a striking reduction or complete absence of Dach1-positive podocytes, respectively (Fig. 8D). In severely affected glomeruli, the nephrin staining was essential to confirm the identification of actual glomeruli, as Dach1-positive nuclei were entirely absent in some cases. Unbiased quantification of Dach1-positive cells from 30 glomeruli (three young and three aged mice) revealed a significant reduction in differentiated Dach1-positive podocytes in 18-month-old mice compared to 6-month-old controls (Fig. 8E).

**Figure 8:**
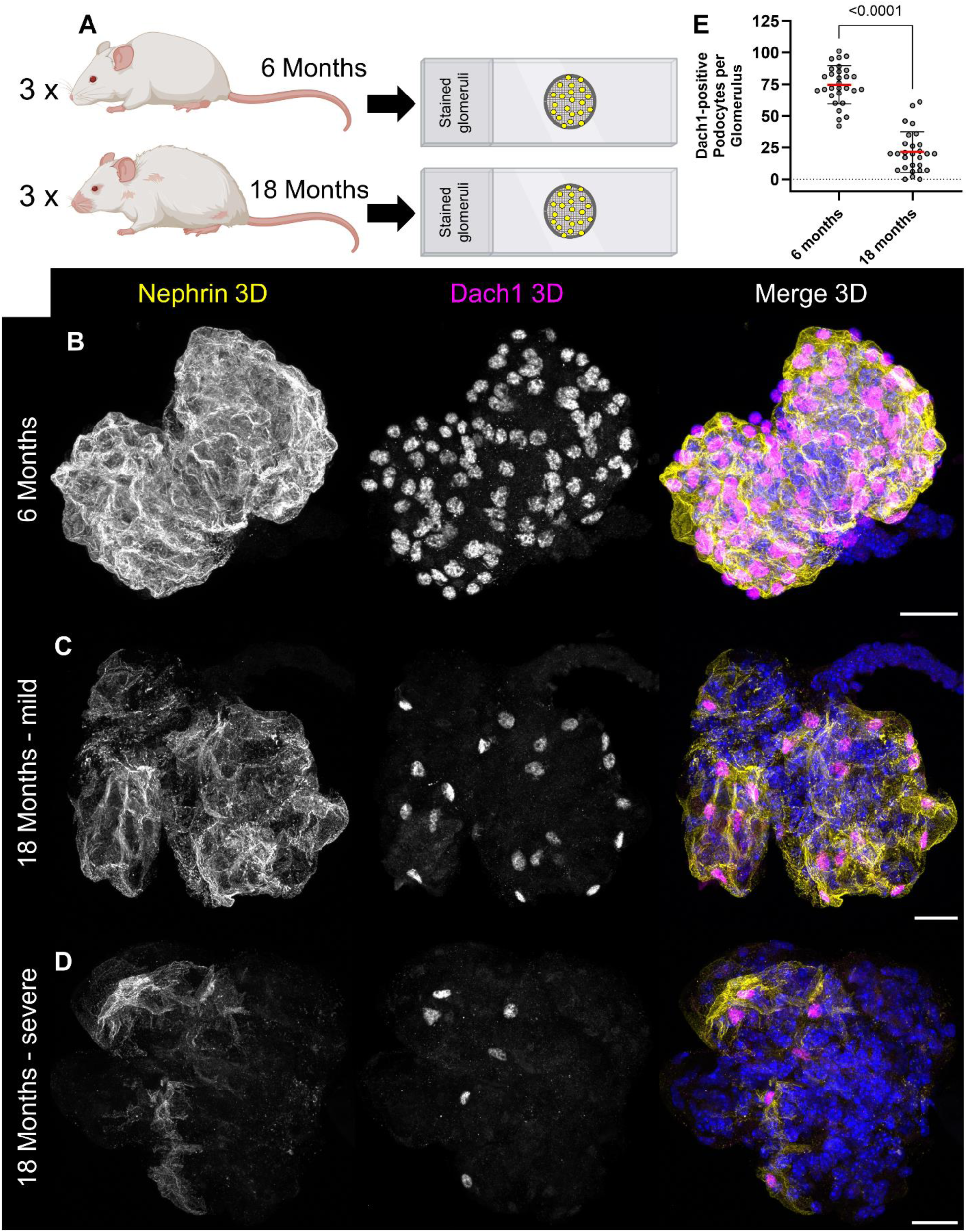
Immunofluorescence of whole glomeruli and podocyte counts in young and aged mice. Glomeruli from 6-month-old (young) and 18-month-old (aged) mice were isolated and stained for Nephrin and Dach1 using the batch immunofluorescence protocol (**A**). MIPs of glomeruli from young mice showed a continuous, linear Nephrin pattern (yellow) and a high density of Dach1-positive podocyte nuclei (magenta, **B**). Glomeruli form aged mice exhibited either a mild phenotype, with partially disrupted Nephrin labeling and a reduced podocyte density (**C**), or a severe phenotype, with marked structural deterioration and near-complete loss of Nephrin and Dach1 signals (**D**). Quantification of Dach1-positive nuclei revealed a significant reduction in podocyte number in aged mice (mean: 21.4, SD: 16.2, n=28) compared to young controls (mean: 74.6, SD: 15.2, n=29 (**E**)). Data were analyzed by a two-tailed Mann-Whitney test; p ≤ 0.05 was considered significant. Scale bars: 20 µm.

These results demonstrate that the *Glomage* podocyte count can be effectively transferred to mammalian glomeruli, enabling rapid and reliable determination of total podocyte number.

## Discussion

Podocytes are crucial cells for maintaining the integrity of the filtration barrier in the kidney.^39^ Damage or loss of podocytes leads to a breakdown of this barrier, allowing high molecular weight proteins such as albumin to pass into the urine – a life-shortening and life-threating situation.^40^ In approximately 75% of patients with chronic kidney disease, postmitotic podocytes are affected, underscoring their critical role and vulnerability.^41^ This circumstance makes it necessary to explore new approaches in drug discovery and analysis. Zebrafish larvae play an important role in this process, as they represent an outstanding model that enables rapid and reliable investigation of kidney development, physiological function, and drug identification.^42–44^ Additionally, the zebrafish is becoming an increasingly important model, as there is a global trend away from mammalian experiments toward simpler organisms and organoids.

Since zebrafish have a high reproduction rate and the larvae are optical transparent, they are ideally suited for *in vivo* high-throughput screening approaches.^16,45,46^ Nevertheless, the subsequent histological analysis of zebrafish larvae remains a significant and time-consuming challenge that has not yet been solved until now.

In the present study, we introduce *Glomage* as a novel high-throughput histological technique designed to accelerate and improve glomerular research. The method is based on batch staining of isolated zebrafish glomeruli, and we further demonstrate that it can also be applied to mice, making it a highly versatile.

Currently, entire zebrafish larvae, which are only a few millimeters in length, are processed individually for histological analysis following experimental procedures. Due to the loss of endogenous fluorophores during paraffin embedding, transgenic larvae are preferably prepared as cryosections, a technically demanding and time-consuming process.

Furthermore, each section must be cut precisely through the single glomerulus of each larva. To overcome these limitations, we developed a method, which allows for a significantly faster and more efficient histological workflow.

The *Glomage* procedure reliably releases whole intact glomeruli from a large number of pre-fixed larvae in a short period of time, it takes only one hour for 100 larvae. In contrast, a skilled operator embeds, orientates and generates cryosections from four larvae in one hour. Therefore, the *Glomage* sample preparation is 25-fold faster and less technically demanding. Additionally, because the samples require no cryoprotection and are not prone to freezing-induced morphological changes, the protocol better preserves its native state.^47^

Another pioneering step of the *Glomage* method is the incubation with antibodies in mesh baskets containing large numbers of glomeruli. In standard stainings of cryosections, 100 µl of antibody solution is required per larva. A theoretical staining of 50 sections would thus requires around 5 ml of antibody solutions. In contrast, the *Glomage* approach allows the simultaneous staining of over 50 glomeruli using only 500 µl of antibody solution, followed by mounting on a single glass slide, highlighting the protocol’s remarkable efficiency. Overall, processing 100 larvae yielded approximately 70 stained and mounted glomeruli within one and a half days, demonstrating and scalability of the workflow.

An important feature of the glomerulus is its three-dimensional architecture, which cannot be fully represented in standard histological sections. To date, analyses of whole glomeruli of larval zebrafish rely on either *in vivo* multi-photon microscopy or whole-mount immunostainings.^36,48–51^ Although these techniques are powerful tools, they have notable limitations. *In vivo* multiphoton microscopy requires specialized equipment and skilled operators, and it does not allow antibody-based detection of specific proteins. Whole-mount immunofluorescence is labor-intensive, often affected by limited antibody penetration and background fluorescence, and neither method is easily scalable.

Here we show that by using antibodies against key podocyte-specific proteins such as nephrin and podocin, we were able to reconstruct the entire slit diaphragm of larval zebrafish glomeruli in three-dimensions. Staining for Pax2, a transcription factor specific for parietal epithelial cells, allows the three-dimensional visualization of the Bowman’s capsule.

Beside this, the determination of the number of podocytes in glomeruli is a highly valuable diagnostic parameter for kidney health and disease. ^52–54^ Although the number of podocytes has already been determined in mouse glomeruli, the corresponding number for the larval zebrafish model was previously unknown. To address this, our protocol exploits the free nuclear diffusion of endogenous fluorescent reporters such as eGFP or mCherry, making it highly specific for podocytes and antibody-independent. By combining the *Glomage* workflow with AI-assisted, unbiased segmentation of eGFP-positive nuclei in our podocyte reporter strains, we can now quantify the number of podocytes rapidly and reproducibly. Using this approach, we quantified podocyte numbers in larvae at 5 dpf. On average, these larvae possess 90-100 podocytes, closely matching the podocyte count observed in healthy adult mouse glomeruli.^9,10^

For the first time, this technique allows quickly longitudinal analysis of podocyte number dynamics during development or disease, and it can be readily adapted to any glomerular cell type with a suitable nuclear marker or reporter.

As a proof-of-concept of this strategy, we used the podocyte counting pipeline for our established FSGS-like model in zebrafish larvae. For this, *Cherry* larvae were exposed to the prodrug NFP which induces a podocyte-specific injury, resulting in many hallmarks of FSGS such as podocyte loss, foot process effacement, glomerular matrix accumulation, proteinuria, and edema formation.^16,29,30^ After 24 hours of NFP treatment, we counted the number of podocytes in healthy, diseased and diseased larvae that received pharmacological treatment. Larval FSGS-induction resulted in a significant loss of viable podocytes and we found that concurrent treatment of the NFP-treated larvae with the compound NM_187 completely prevented podocyte loss.

Furthermore, *Glomage* enables volumetric analysis of the mCherry-or eGFP-positive volume in larval glomeruli. In accordance with podocyte loss, NFP induced a significant reduction and the co-treatment with NM_187 a complete rescue of the mCherry-positive volume. This novel readout might be additionally used in the future to assess podocyte hypertrophy or further volumetric changes upon injury or drug treatment.

Beside the use of *Glomage* for imaging purposes, we developed a protocol for the extraction of glomerular RNA. Compared to a whole-larvae RNA isolation, this approach yielded in a strong enrichment of glomerular and podocyte-specific transcripts, while markers of other organs such as pancreas, heart, or eyes were undetectable. This indicates a high purity of glomerular RNA.

With this modified technique, we measured the downregulation of podocyte-specific genes upon podocyte injury as well as a protective effect of NM_187 on mRNA levels. The close agreement between imaging and mRNA results demonstrates that both approaches can be effectively combined, allowing podocyte imaging and RNA analysis from the same batch of larvae which could also be used for RNA_Seq and other omic technologies.

To study whether *Glomage* has also the potential for other animal species, we next applied it to isolated glomeruli from mouse tissue. First, glomeruli were obtained by a sieving and differential adhesion technique from a *nephrin*:CFP reporter strain.^38^ After this, all isolated glomeruli were processed with the same antibody staining procedure in mesh baskets as reported for zebrafish glomeruli. Here, we have shown by using an antibody against the slit-diaphragm protein nephrin, that the *Glomage* workflow is directly transferable to mouse tissue.

As proof-of-concept, we used the technique also in the context of age-associated podocyte loss. To address this this, we isolated and analyzed glomeruli from young and aged mice by using the transcription factor Dach1 as a podocyte-specific nuclear marker. We found that aged mice exhibited a significant reduction in Dach1-positive podocyte number compared with young animals, consistent with previous reports and supporting the validity of this method.^9,10^

In summary, compared to already established protocols, *Glomage* applied to mouse tissue offers several advantages: the entire workflow from living animal to stained and mounted glomeruli can be completed within two days. No animal perfusion, tissue clearing, or prolonged antibody incubation on thick slices is required, steps that typically take up to two weeks.^9^ Furthermore, the method does not require specialized microscopy techniques such as light-sheet or super-resolution imaging. Additionally, the number of glomeruli obtained from one kidney is sufficient for multiple staining procedures, allowing the contralateral kidney to be used for standard histology and omics analysis. Moreover, this approach can be extended to established glomerular injury models such as nephrotoxic serum nephritis or puromycin aminonucleoside nephritis.

In summary, the *Glomage* workflow combines accuracy, scalability, and cross-species applicability with remarkable simplicity and velocity using only standard laboratory equipment. This accessibility allows a wide range of laboratories to implement high-throughput, three-dimensional analyses of glomeruli, supporting studies from basic podocyte biology to translational drug screening, and enabling rapid reproducible insights into kidney health and disease.

## Methods

### Zebrafish husbandry

Zebrafish were maintained under standard conditions as previously described.^55^ The transparent *nphs2*:eGFP (*Podo:GFP*) line expresses eGFP specifically in pronephric podocytes and was generated by crossing TG(*nphs2*:GAL4-VP16) (own outcross) with TG(*UAS*:eGFP) (a kind gift from Prof. Brand, Dresden), resulting in the genotype TG(*nphs2*:GAL4-VP16);TG(*UAS*:eGFP) mitfa^W2/W2^ (Fig. S1, A–C). The *nphs2*:NTR-mCherry (*Cherry*) line expresses both mCherry and the nitroreductase in podocytes and was used for experiments involving targeted podocyte injury. The genotype of this line is TG(*nphs2*:GAL4-VP16);TG(*UAS*:Eco.nfsB-mCherry) mitfa ^W2/W2^ (a kind gift from Prof. Weibin Zhou, Fig. S1, D–E).^56,57^ Fluorophore expression was verified at 4 dpf prior to the start of each experiment under 0.1 mg/ml tricaine anesthesia (MS-222, E10521, Merck, Darmstadt, Germany).

All experiments with zebrafish larvae were terminated at 5 dpf and conducted in accordance with the guidelines of the local regulatory authorities.

### Isolation and collection of larval zebrafish glomeruli

100 *Podo:GFP* larvae were fixed in 2% PFA for 1.5 hours at room temperature. After washout, larvae were transferred in PBS to a screw cap tube prepared with 100 µl ceramic beads. Both the beads and the inner surface of the screw cap were pre-coated with FBS (Thermo Fisher Scientific, Dreieich, Germany). PBS, ceramic beads, and the 100 larvae were adjusted to a total volume of 1 ml and dissociated using a dissociator (Fast-Prep-24^TM^, MP Biomedicals, Eschwege, Germany) at 4 m/s for 20 s. The resulting suspension was then transferred, along with additional PBS to a FBS-coated petri dish (Greiner Bio-One, Kremsmünster, Austria). Fluorescent and morphologically intact glomeruli were manually picked under a fluorescence stereomicroscope (SMZ18, Nikon, Düsseldorf, Germany) using FBS-coated gel-loading pipette tips (Biozym, Hessisch-Oldendorf, Germany) and transferred to a second FBS-coated petri dish containing PBS. After a second round of quality control, glomeruli were transferred to FBS-coated cell culture inserts (ThinCert®, Greiner Bio-One, Cat. No. 662638) with a mesh bottom (pore size: 8 µm) placed in a 24-well plate (Greiner Bio-One).

### Immunofluorescence of whole larval zebrafish glomeruli

All solutions were applied by gently rinsing through the mesh, allowing liquids to pass while retaining the glomeruli. Glomeruli resting on the mesh were washed twice with PBS containing 0.1% Triton X-100 (PBST). Permeabilization was carried out by incubating the samples in 0.3% PBST for 5 minutes, followed by antigen retrieval in 1× sodium dodecyl sulfate (Biomol, Hamburg, Germany) buffer in PBS for 5 minutes. After washing with PBST, blocking solution was applied for 45 minutes. Primary antibodies were diluted in a total volume of 500 µl and incubated overnight at 4°C. The following rabbit primary antibodies and dilutions were used: anti-zebrafish nephrin 1:2000 (Innovagen, Lund, Sweden), anti-podocin 1:7500 (Sigma-Aldrich, Taufkirchen, Germany, Cat. No. PO372), anti-Pax2 1:1000 (Abcam, Cambridge, UK, Cat. No. ab38738), anti-Ehd3 1:200 (Sigma-Aldrich, Cat. No. HPA049986), and anti-Laminin 1:100 (Sigma-Aldrich, Cat. No. L9393). After antibody incubation, glomeruli were washed four times for 5 minutes each with PBST on a horizontal shaker. This was followed by a 1-hour incubation with an Alexa Fluor 647-conjugated donkey anti-rabbit secondary antibody (Thermo Fisher Scientific) at a dilution of 1:800. Glomeruli were subsequently washed four times with PBST. Nuclei were counterstained using HOECHST 33342 (Sigma-Aldrich), and F-actin was labeled with Phalloidin-546, applied together with the nuclear stain. After two additional washes with PBST and three final washes with A. dest., the mesh was carefully cut using a scalpel, grasped with forceps, and mounted on a glass slide containing a drop of Mowiol (Roth) for embedding with a 22 x 22 mm cover slit.

### Imaging of whole zebrafish glomeruli

All glomeruli were imaged using an FV3000 confocal microscope (Evident, Tokyo, Japan) equipped with a 1.25x air objective (NA: 0.04), a 20x air objective (NA: 0.8), and a 60x oil immersion objective (NA: 1.5). The 1.25x objective was used to image the entire mesh for the generation of a map, using brightfield and either 488 nm or 561 nm excitation depending on the expressed fluorophore (488 nm for eGFP, 561 nm for mCherry). This map enabled automated navigation of the 20x and 60x objective to individual glomeruli. For each glomerulus, z-stacks were acquired with the 60x objective and a step size of 1 µm. The zoom factor used for acquisition ranged from 2x to 3.5x, depending on the individual orientation of the zebrafish glomerulus in Mowiol.

### Cryosections and immunofluorescence

Zebrafish larvae (*Podo:GFP*) were fixed in 2% PFA at 5 dpf for 1.5 hours at room temperature. Larvae were incubated overnight in 15% sucrose (Sigma-Aldrich) in PBS after fixation. Following this, larvae were snap frozen in a 1:1 solution of 15% sucrose in PBS and TissueTek O.C.T. compound (Sakura Finetek Europe, AV, Netherlands) in liquid nitrogen. 5 µm cryosections were prepared using a CM 1950 cryostat (Thermo Fisher Scientific) and stained with the same primary and secondary antibodies at previously described concentrations. Sections were mounted in Mowiol and images were captured with the Evident FV3000 system using a 60x water objective (NA: 1.2).

### Image processing

All imaging data were processed using the open-source software FIJI.^58^ Custom-written macros were employed to automatically generate single slices and maximum intensity projections (MIPs) from z-stacks. The number of glomeruli per mesh was determined manually. Two-channel hyperstacks (nuclei + eGFP or nuclei + mCherry) were used for glomerular reconstruction, quantification of eGFP- or mCherry-positive podocyte nuclei and nuclear volume using Imaris (Bitplane, version 10.2.0). Nuclei were segmented using a custom trained AI network based on manually labelled nuclei (Fig. 3) or threshold segmentation (Fig. 4, 5, 8) followed by watershed-based separation of touching objects (region growing, seed size 2 µm) and filtering for total volume. eGFP or mCherry positive nuclei were identified by an intensity mean threshold filter. The podocyte volume was determined by threshold segmentation and volume filtering.

### Induction of larval FSGS and compound treatment

Specific podocyte injury was induced by treating 100 *Cherry* larvae with 50 nM nifurpirinol (NFP, Sigma-Aldrich) in 0.5x E3 medium for 24 hours from 4 to 5 dpf. The healthy control group received 0.2% DMSO, whereas the intervention group was treated with 50 nM NFP in combination with the compound NM_187 for the same duration. NM_187 is an experimental small molecule. Its structure and mechanism of action are currently not disclosed due to intellectual property restrictions. Larvae were subsequently used for glomerular isolation, immunofluorescence, and confocal imaging (as described above), or processed fresh for RNA isolation.

### RNA isolation and PCR of zebrafish glomeruli

Non-fixed *Cherry* zebrafish larvae were dissociated as described above at 4 m/s for 1 s. Glomeruli were collected under fluorescence guidance, and RNA was isolated following two rounds of quality control using the Single Cell RNA Purification Kit (Norgen Biotek, Thorold, ON, Canada) after manufacturer‘s instructions. RNA from 10 whole larvae, processed in parallel, served as controls. Reverse transcription was carried out with the QuantiTect® Reverse Transcription Kit (Qiagen, Hilden, Germany), followed by RT–PCR using DreamTaq DNA Polymerase (Thermo Fisher Scientific) for 34 cycles. RT-qPCR was performed as previously described on a QuantStudio™ 3 Real-Time PCR System (Thermo Fisher Scientific) using the iTaq Universal SYBR Green Supermix (Bio-Rad).^59^ Raw Ct-values were normalized to *eef1a1* as reference gene and to whole larvae or to the 0.2% DMSO treatment group, as indicated respectively. Ct-values ≤ 38 were excluded from analysis. As negative controls we included no-template controls (NTC) as well as no-reverse-transcriptase controls (-RT). All measurements were run in triplicates and primer sequences are provided in the supplemental material (Table S1).

### Mouse husbandry

For this study, we used transgenic *nephrin*:CFP mice, in which CFP is expressed specifically in podocytes under the control of a *nephrin* promoter fragment. The generation of this mouse line involved insertion of the CFP cDNA into the EcoRI site of the NPXRS *nephrin* construct, as previously described.^60,61^ Animals were housed under standardized conditions (21 °C, 60% humidity, 12:12 hours light–dark cycle) with ad libitum access to food and water. Experimental procedures were carried out on 6-month-old or 18-month-old mice. All animal work was conducted in compliance with national animal welfare legislation with prior approval from the competent local authority.

### Isolation and immunofluorescence of mouse glomeruli

Glomeruli were isolated using the differential adhesion method as described by Wang et al.. ^38^ At the final step, glomeruli were fixed in 2% PFA for 20 minutes at room temperature in a 15 ml conical tube (Sarstedt, Nümbrecht, Germany). After fixation, PFA was removed by washing, and all glomeruli from one mouse were resuspended in 2 ml PBS. 50 µl of the suspension was inspected under a stereomicroscope (brightfield) to assess the density of glomeruli in the solution. An appropriate volume of the suspension was then transferred onto cell culture inserts (Greiner Bio-One) to achieve an estimated number of ≥50 glomeruli per mesh. Immunofluorescence staining, embedding, and imaging were performed identically to the procedure used for isolated zebrafish glomeruli. The following antibodies and concentrations were used for the immunofluorescence on mouse glomeruli: Anti-Nephrin 1:1000 (Progen, Heidelberg, Germany, Cat. No.: GP-N2), Anti-Dach1 1:500 (Sigma-Aldrich, Cat. No.: HPA012672). The same Alexa Fluor 647-conjugated donkey anti-rabbit (Thermo Fisher Scientific) and a Cy3-conjugated donkey-anti guinea pig (Jackson ImmunoResearch, Newmarket, United Kingdom) were used as secondary antibodies. 4′,6-diamidino-2-phenylindole (DAPI, Sigma-Aldrich) was used for the nuclear counterstain.

### Statistical analysis

Data analysis and graph generation were performed using GraphPad Prism 9.1.2 (GraphPad Software, San Diego, CA, USA). Normality was assessed using the Shapiro-Wilk test. For comparisons between two groups, either an unpaired two-tailed Student’s t-test or a Mann-Whitney U test was applied. For comparisons between more than two groups, a one-way ANOVA followed by Tukey’s multiple comparisons test or a Kruskal-Wallis test with Dunn’s post hoc correction was used, as appropriate. A p-value ≤0.05 was considered statistically significant.

## Supporting information

Supplemental Material Glomage

## Acknowledgements

The authors are very grateful for excellent technical assistance of Jurij Barmenkov, Claudia Weber and the outstanding zebrafish as well as mouse husbandry by Oliver Zabel and Steffen Prellwitz. Image analysis of S.S.L. was conducted at the Center for Microscopy and Image Analysis of the University of Zurich. Selected elements of Figures 1, 6, 7, and 8 were created using BioRender.com.

## Author contributions

Conceptualization: M.S., N.E.

Methodology: M.S., S.M.B., T.L.

Software: M.S., T.L., S.S.L.

Validation: M.S., T.L.

Formal analysis: M.S., T.L., S.S.L.

Investigation: M.S., T.L., S.M.B.

Resources: S.S.L., N.E.

Data curation: M.S.,T.L., S.S.L.

Writing – original draft: M.S.

Writing – review & editing; T.L., S.S.L. N.E.

Visualization: M.S., S.S.L.

Supervision: N.E.

Project administration: N.E.

Funding acquisition: N.E.

## Data and materials available

All data needed to evaluate the conclusions are present in the manuscript and/or the Supplemental Materials.

## Supplementary Material

The corresponding supplementary information can be found in the PDF file:

**Supplemental Material Glomage**

**Other Supplementary Material for this manuscript includes the following:**

Movie S1: https://doi.org/10.6084/m9.figshare.30384343

Movie S2: https://doi.org/10.6084/m9.figshare.30384454

Movie S3: https://doi.org/10.6084/m9.figshare.30384496

Movie S4: https://doi.org/10.6084/m9.figshare.30384502

Movie S5: https://doi.org/10.6084/m9.figshare.30384526

Movie S6: https://doi.org/10.6084/m9.figshare.30425632

## References

1. Bikbov, B. et al. Global, regional, and national burden of chronic kidney disease, 1990–2017: a systematic analysis for the Global Burden of Disease Study 2017. The Lancet 395, 709–733 (2020).

2. Foreman, K. J. et al. Forecasting life expectancy, years of life lost, and all-cause and cause-specific mortality for 250 causes of death: reference and alternative scenarios for 2016-40 for 195 countries and territories. Lancet 392, 2052–2090 (2018).

3. Pavenstädt, H., Kriz, W. & Kretzler, M. Cell Biology of the Glomerular Podocyte. Physiological Reviews 83, 253–307 (2003).

4. Miyaki, T. et al. Three-dimensional imaging of podocyte ultrastructure using FE-SEM and FIB-SEM tomography. Cell Tissue Res 379, 245–254 (2020).

5. Kocylowski, M. K. et al. A slit-diaphragm-associated protein network for dynamic control of renal filtration. Nat Commun 13, 6446 (2022).

6. Bai, X. Y. & Basgen, J. M. Podocyte Number in the Maturing Rat Kidney. Am J Nephrol 33, 91–96 (2011).

7. White, K. E. & Bilous, R. W. Estimation of podocyte number: A comparison of methods. Kidney International 66, 663–667 (2004).

8. Puelles, V. G. et al. Design-based stereological methods for estimating numbers of glomerular podocytes. Annals of Anatomy-Anatomischer Anzeiger 196, 48–56 (2014).

9. Puelles, V. G. et al. Validation of a Three-Dimensional Method for Counting and Sizing Podocytes in Whole Glomeruli. J Am Soc Nephrol 27, 3093–3104 (2016).

10. Shi, J. et al. Quantifying Podocyte Number in a Small Sample Size of Glomeruli with CUBIC to Evaluate Podocyte Depletion of db/db Mice. Journal of Diabetes Research 2023, 1–12 (2023).

11. Puelles, V. G., Combes, A. N. & Bertram, J. F. Clearly imaging and quantifying the kidney in 3D. Kidney International 100, 780–786 (2021).

12. Drummond, I. A. et al. Early development of the zebrafish pronephros and analysis of mutations affecting pronephric function. Development 125, 4655–4667 (1998).

13. Drummond, I. A. & Davidson, A. J. Zebrafish Kidney Development. in Methods in Cell Biology vol. 100 233–260 (Elsevier, 2010).

14. Schindler, M. & Endlich, N. Zebrafish as a model for podocyte research. American Journal of Physiology-Renal Physiology 326, F369–F381 (2024).

15. Howe, K. et al. The zebrafish reference genome sequence and its relationship to the human genome. Nature 496, 498–503 (2013).

16. Schindler, M. et al. A Novel High-Content Screening Assay Identified Belinostat as Protective in a FSGS-Like Zebrafish Model. J Am Soc Nephrol 34, 1977–1990 (2023).

17. Schindler, M., Blumenthal, A., Moeller, M. J., Endlich, K. & Endlich, N. Adriamycin does not damage podocytes of zebrafish larvae. PLoS ONE 15, e0242436 (2020).

18. Yu, T. et al. Identification of renal stem cells in zebrafish. Sci Adv 11, eadx5296 (2025).

19. Müller, T. et al. Non-muscle myosin IIA is required for the development of the zebrafish glomerulus. Kidney International 80, 1055–1063 (2011).

20. Bolten, J. S., Pratsinis, A., Alter, C. L., Fricker, G. & Huwyler, J. Zebrafish ( *Danio rerio*) larva as an in vivo vertebrate model to study renal function. American Journal of Physiology-Renal Physiology 322, F280–F294 (2022).

21. Siegerist, F. et al. The differential expression of MAGI2 in glomerulopathies and its application as a molecular discriminator of podocytopathies. J Transl Med 23, 701 (2025).

22. Bondue, T. et al. Evaluation of the efficacy of cystinosin supplementation through CTNS mRNA delivery in experimental models for cystinosis. Sci Rep 13, 20961 (2023).

23. Taimatsu, K. et al. Comprehensive 3D Imaging of Whole Zebrafish Using a Water-Based Clearing Reagent for Hard Tissues. Zebrafish 22, 65–75 (2025).

24. Siegerist, F. et al. Evaluation of endogenous miRNA reference genes across different zebrafish strains, developmental stages and kidney disease models. Sci Rep 11, 22894 (2021).

25. Mattias, F. et al. Regulation of the transcriptome, miRNAs, and alternative splicing in a FSGS zebrafish injury model. Preprint at 10.1101/2025.07.09.663814 (2025).

26. Arber, D. A. Effect of Prolonged Formalin Fixation on the Immunohistochemical Reactivity of Breast Markers: Applied Immunohistochemistry & Molecular Morphology 10, 183–186 (2002).

27. Konno, K., Yamasaki, M., Miyazaki, T. & Watanabe, M. Glyoxal fixation: An approach to solve immunohistochemical problem in neuroscience research. Sci Adv 9, eadf7084 (2023).

28. Dekan, G., Gabel, C. & Farquhar, M. G. Sulfate contributes to the negative charge of podocalyxin, the major sialoglycoprotein of the glomerular filtration slits. Proc. Natl. Acad. Sci. U.S.A. 88, 5398–5402 (1991).

29. Kerjaschki, D. Caught flat-footed: podocyte damage and the molecular bases of focal glomerulosclerosis. J Clin Invest 108, 1583–1587 (2001).

30. Klawitter, M. et al. Investigating FSGS-like injury in zebrafish larvae by nifurpirinol: efficacy and molecular insight. Am J Physiol Renal Physiol 327, F463–F475 (2024).

31. Hansen, K. U. I. et al. Prolonged podocyte depletion in larval zebrafish resembles mammalian focal and segmental glomerulosclerosis. FASEB j. 34, 15961–15974 (2020).

32. George, M. et al. Renal thrombotic microangiopathy in mice with combined deletion of endocytic recycling regulators EHD3 and EHD4. PLoS One 6, e17838 (2011).

33. Wanner, N. et al. Unraveling the role of podocyte turnover in glomerular aging and injury. J Am Soc Nephrol 25, 707–716 (2014).

34. Wan, X. et al. Loss of Epithelial Membrane Protein 2 Aggravates Podocyte Injury via Upregulation of Caveolin-1. Journal of the American Society of Nephrology 27, 1066–1075 (2016).

35. Siegerist, F., Blumenthal, A., Zhou, W., Endlich, K. & Endlich, N. Acute podocyte injury is not a stimulus for podocytes to migrate along the glomerular basement membrane in zebrafish larvae. Sci Rep 7, 43655 (2017).

36. Siegerist, F., Zhou, W., Endlich, K. & Endlich, N. 4D in vivo imaging of glomerular barrier function in a zebrafish podocyte injury model. Acta Physiol (Oxf) 220, 167–173 (2017).

37. Kindt, F. et al. A novel assay to assess the effect of pharmaceutical compounds on the differentiation of podocytes. Br J Pharmacol 174, 163–176 (2017).

38. Wang, H. et al. A simple and highly purified method for isolation of glomeruli from the mouse kidney. Am J Physiol Renal Physiol 317, F1217–F1223 (2019).

39. Loreth, D., Sachs, W. & Meyer-Schwesinger, C. The Life of a Kidney Podocyte. Acta Physiologica 241, e70081 (2025).

40. Nagata, M. Podocyte injury and its consequences. Kidney International 89, 1221–1230 (2016).

41. Wiggins, R.-C. The spectrum of podocytopathies: A unifying view of glomerular diseases. Kidney International 71, 1205–1214 (2007).

42. Poureetezadi, S. J. & Wingert, R. A. Little fish, big catch: zebrafish as a model for kidney disease. Kidney International 89, 1204–1210 (2016).

43. Kramer-Zucker, A. G., Wiessner, S., Jensen, A. M. & Drummond, I. A. Organization of the pronephric filtration apparatus in zebrafish requires Nephrin, Podocin and the FERM domain protein Mosaic eyes. Developmental Biology 285, 316–329 (2005).

44. Peng, Z. et al. Somites are a source of nephron progenitors in zebrafish. Nat Commun 16, 6914 (2025).

45. Gehrig, J., Pandey, G. & Westhoff, J. H. Zebrafish as a Model for Drug Screening in Genetic Kidney Diseases. Front. Pediatr. 6, 183 (2018).

46. Bolten, J. S. et al. Zebrafish (Danio rerio) larvae as a predictive model to study gentamicin-induced structural alterations of the kidney. PLoS ONE 18, e0284562 (2023).

47. Adam, M., Fei Hu, J., Lange, P. & Wolfinbarger, L. The effect of liquid nitrogen submersion on cryopreserved human heart valves. Cryobiology 27, 605–614 (1990).

48. Endlich, N. et al. Two-photon microscopy reveals stationary podocytes in living zebrafish larvae. J Am Soc Nephrol 25, 681–686 (2014).

49. Siegerist, F., Blumenthal, A., Zhou, W., Endlich, K. & Endlich, N. Acute podocyte injury is not a stimulus for podocytes to migrate along the glomerular basement membrane in zebrafish larvae. Sci Rep 7, 43655 (2017).

50. Ichimura, K. et al. Developmental Localization of Nephrin in Zebrafish and Medaka Pronephric Glomerulus. J Histochem Cytochem. 61, 313–324 (2013).

51. Gerlach, G. F. & Wingert, R. A. Zebrafish pronephros tubulogenesis and epithelial identity maintenance are reliant on the polarity proteins Prkc iota and zeta. Developmental Biology 396, 183–200 (2014).

52. Puelles, V. G. & Bertram, J. F. Counting glomeruli and podocytes: rationale and methodologies. Current Opinion in Nephrology and Hypertension 1 (2015) doi:10.1097/MNH.0000000000000121.

53. Haruhara, K. et al. Podocyte density as a predictor of long-term kidney outcome in obesity-related glomerulopathy. Kidney International 106, 496–507 (2024).

54. Ding, F. et al. Accelerated podocyte detachment and progressive podocyte loss from glomeruli with age in Alport Syndrome. Kidney International 92, 1515–1525 (2017).

55. Endlich, N. et al. The transcription factor Dach1 is essential for podocyte function. J Cell Mol Med 22, 2656–2669 (2018).

56. Wan, X. et al. Loss of Epithelial Membrane Protein 2 Aggravates Podocyte Injury via Upregulation of Caveolin-1. J Am Soc Nephrol 27, 1066–1075 (2016).

57. Zhou, W. & Hildebrandt, F. Inducible Podocyte Injury and Proteinuria in Transgenic Zebrafish. Journal of the American Society of Nephrology 23, 1039–1047 (2012).

58. Schindelin, J., et al. Fiji: an open-source platform for biological-image analysis. Nat Methods 9, 676–682 (2012).

59. Ristov, M.-C. et al. The ShGlomAssay Combines High-Throughput Drug Screening With Downstream Analyses and Reveals the Protective Role of Vitamin D3 and Calcipotriol on Podocytes. Front. Cell Dev. Biol. 10, 838086 (2022).

60. Cui, S., Li, C., Ema, M., Weinstein, J. & Quaggin, S. E. Rapid Isolation of Glomeruli Coupled with Gene Expression Profiling Identifies Downstream Targets in Pod1 Knockout Mice. Journal of the American Society of Nephrology 16, 3247–3255 (2005).

61. Wong, M. A., Cui, S. & Quaggin, S. E. Identification and characterization of a glomerular-specific promoter from the human nephrin gene. American Journal of Physiology-Renal Physiology 279, F1027–F1032 (2000).

