## Supplemental Material Glomage for "*Glomage*: A multimodal platform for high-content morphological and RNA profiling of glomeruli in zebrafish and mouse models"

### Table of contents

#### Supplemental Figures


1. Fig. S1
2. Fig. S2
3. Fig. S3
4. Fig. S4
5. Fig. S5


#### Supplemental Tables


1. Table S1


#### Supplemental Movie Captions


1. Movie S1
2. Movie S2
3. Movie S3
4. Movie S4
5. Movie S5
6. Movie S6

### Supplemental Figures

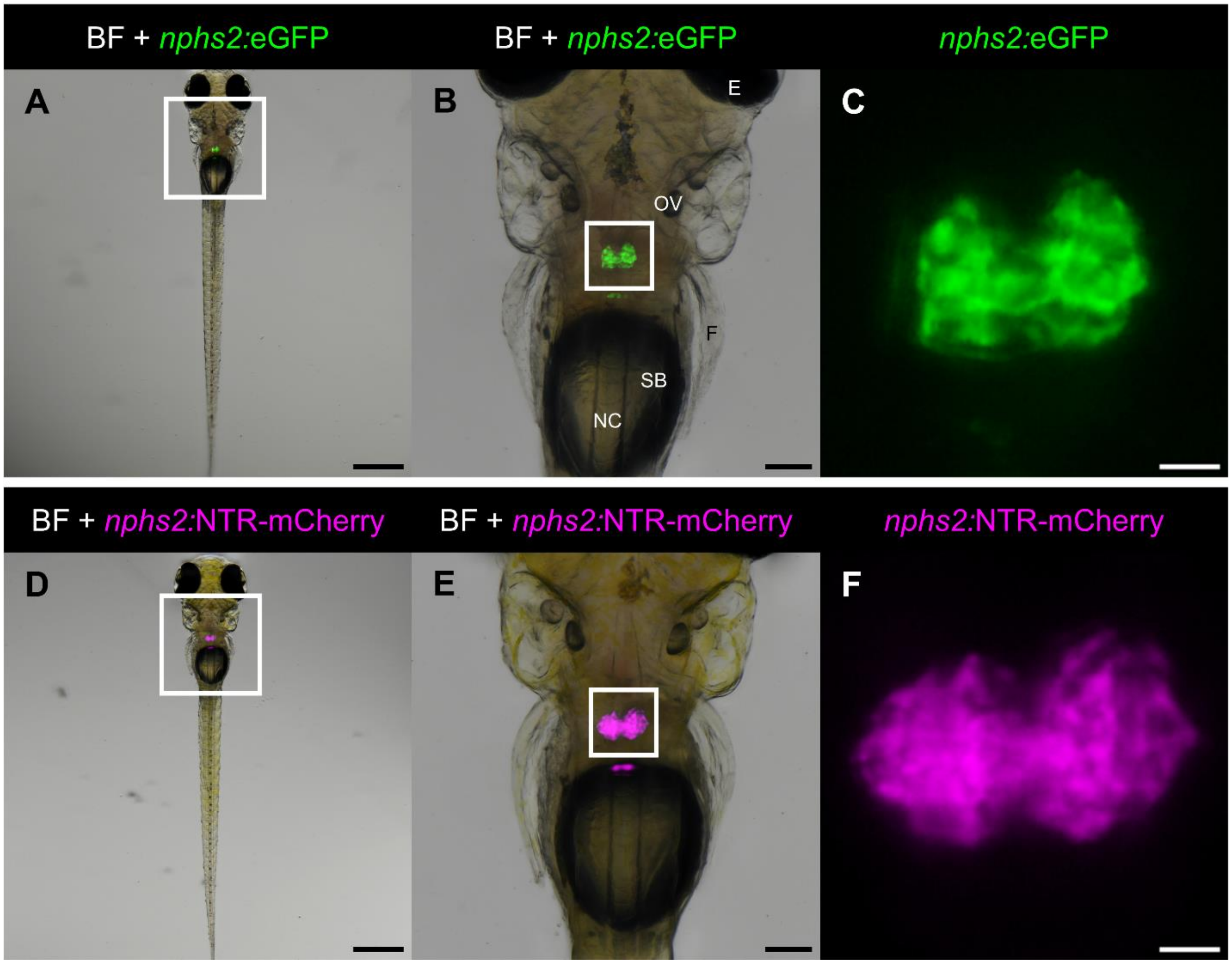


**Figure S1: Overview of the transgenic zebrafish lines and their expression patterns.** Larvae of the *Podo:GFP* line express eGFP exclusively in podocytes, the glomerulus is located cranial to the swim bladder and exhibits the typical peanut-like shape at 5 dpf (A–C). The *Cherry* strain shows the same expression pattern, with the fluorophore mCherry expressed in podocytes. In addition, this strain expresses *E. coli* nitroreductase (NTR) specifically in podocytes (D–F). NTR renders this strain sensitive to podocyte injury upon treatment with nifurpirinol (NFP). BF: Brightfield, E: Eye; OV: Otic vesicle; F: Fin; SB: Swim bladder; NC: Notochord. Scale bars: 500 µm (A, D); 100 µm (B, E); 20 µm (C, F).

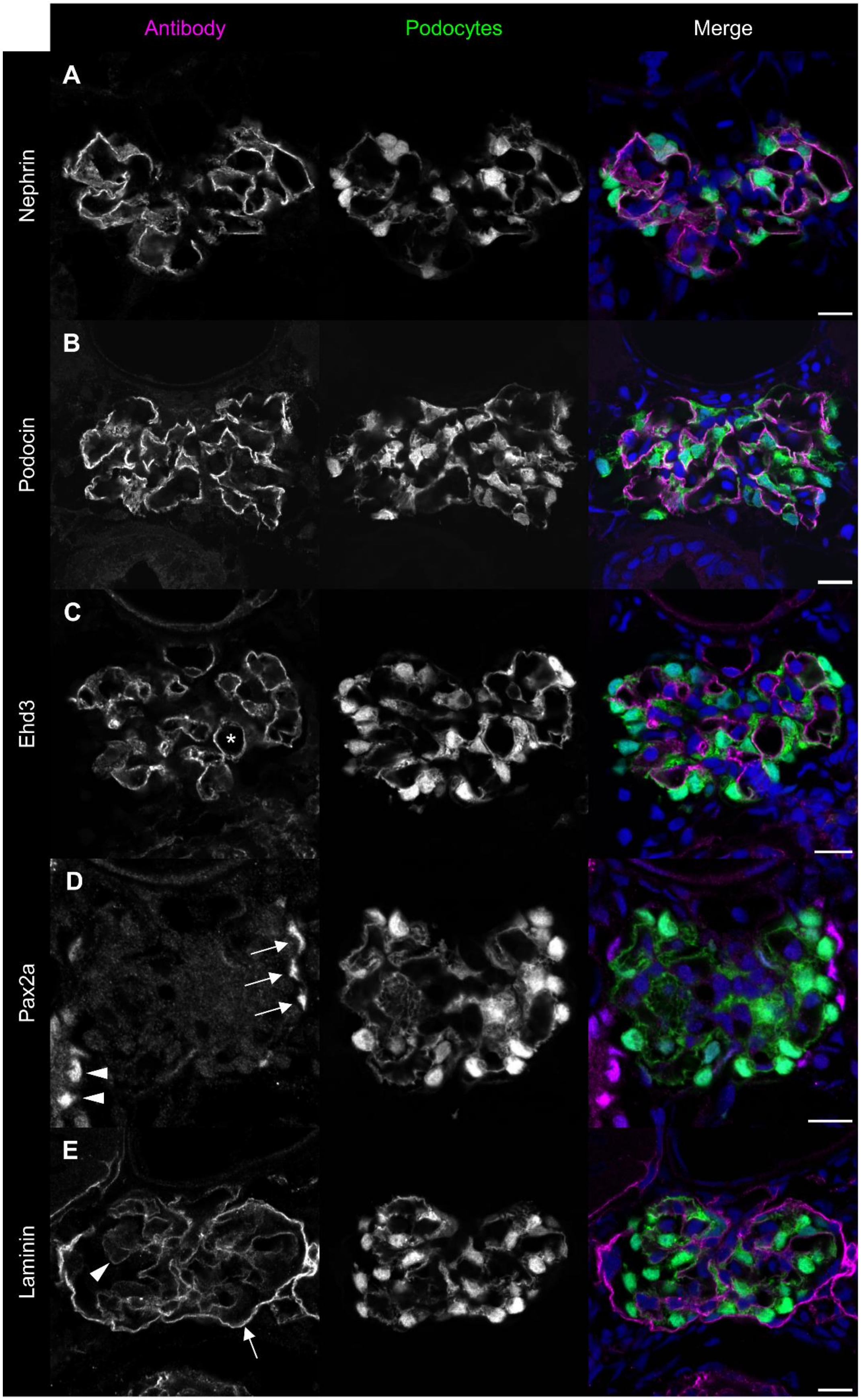

Antibody
Podocytes
Merge
A
Nephrin
B
Podocin
C
Ehd3
*
D
Pax2a
E
Laminin

**Figure S2: Glomerular cryosections of *Podo:GFP* larvae stained with various antibodies.** Nephrin and Podocin label the slit diaphragm of the glomerular filtration barrier. Ehd3 marks fenestrated glomerular endothelial cells revealing cross sections through glomerular capillaries (asterisk). Pax2 labels parietal epithelial cells (arrows) and proximal tubule cells (arrowheads). The Laminin antibody highlights basement membranes, including the parietal (arrow) and glomerular (arrowhead) basement membranes. Scale bars: 10 µm.

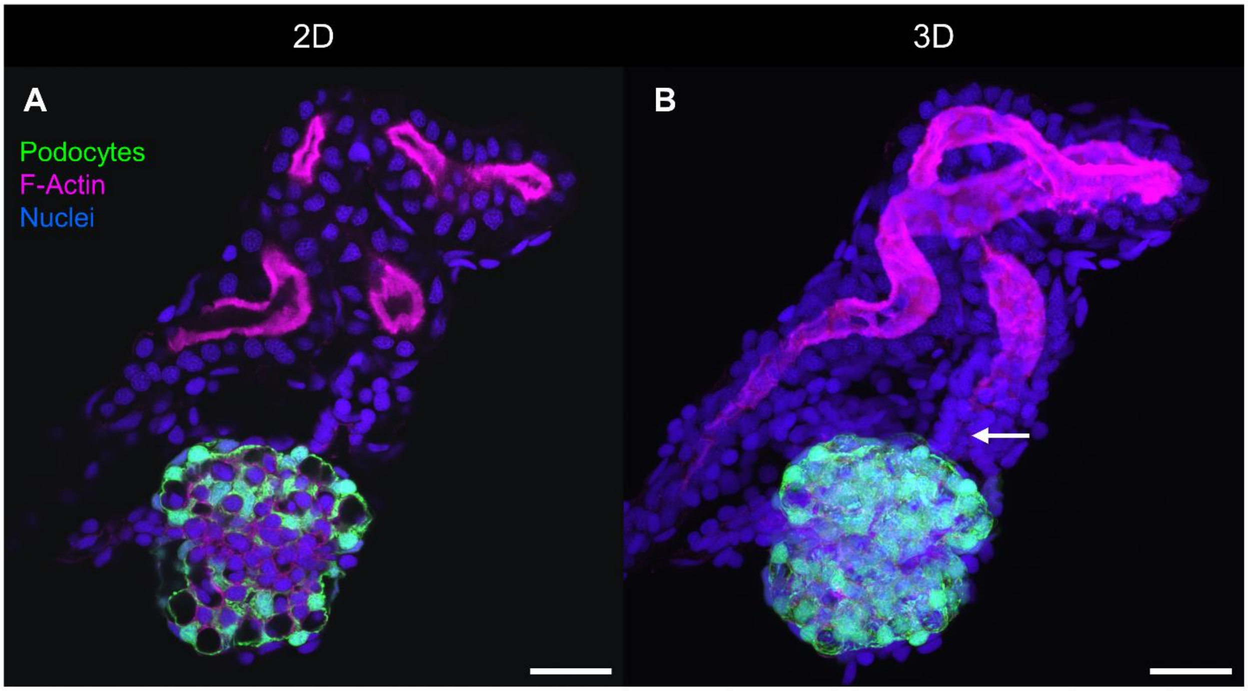


**Figure S3: Two- and three-dimensional visualization of a glomerulus and tubule co-isolation.** Podocytes are shown in green (Podo:GFP), while F-actin, labeled with Phalloidin, marks the brush border of proximal tubule cells (magenta). A single optical section shows different angles of a cut through the convoluted tubule (A). The MIP displays the entire convoluted tubule with its complete brush border (asterisk). The neck segment, which connects the tubule to the glomerulus, lacks a brush border (arrow) (B). Scale bars: 20 µm.

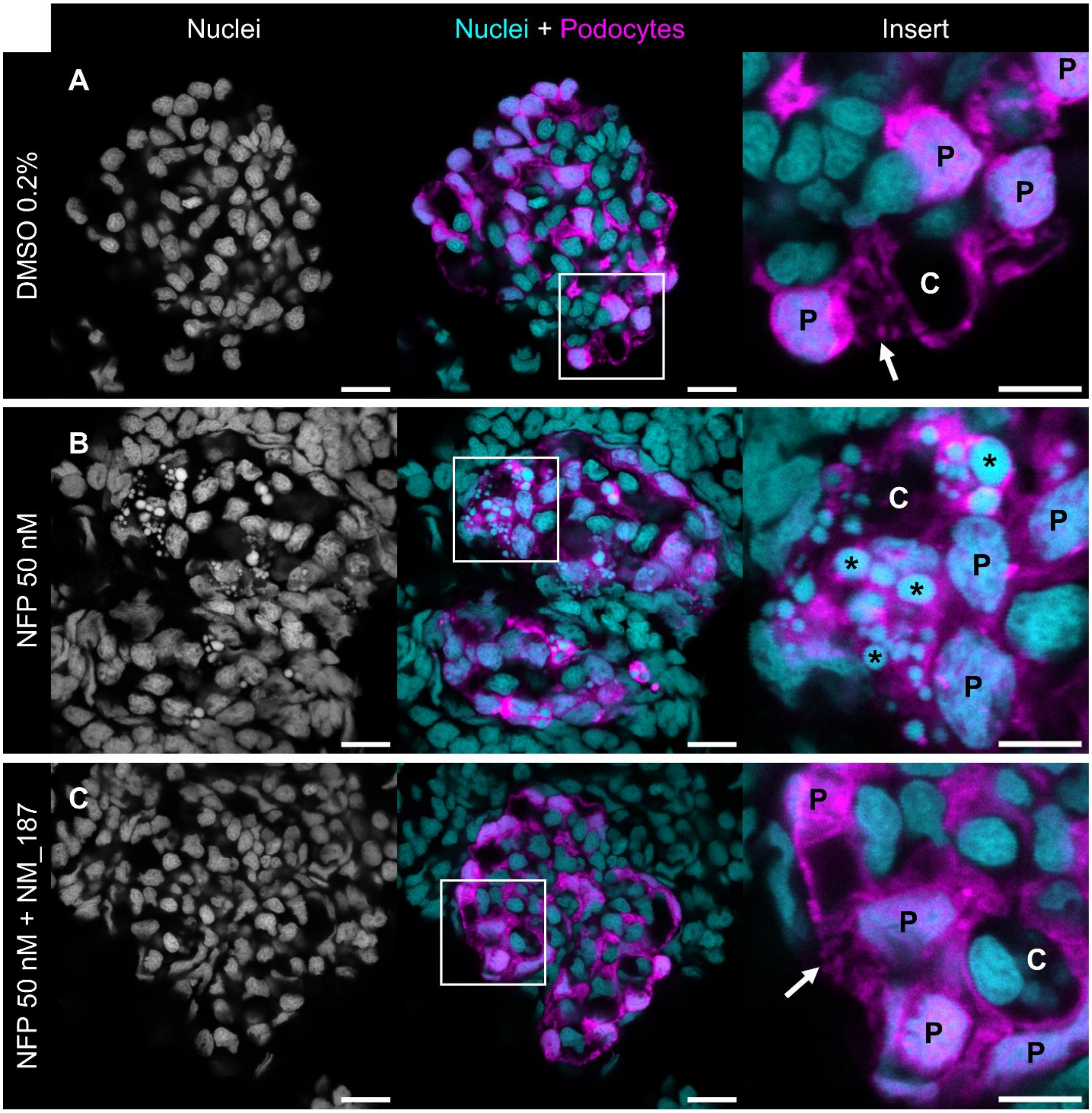


**Figure S4: Single slices of whole glomeruli from *Cherry* larvae treated with DMSO, NFP, and NFP + NM_187.** In the healthy glomerulus (DMSO), nuclei (cyan) appear normal, and primary processes of podocytes (magenta) are visible (arrow) (A). Larvae treated with NFP display fragmented podocyte nuclei (asterisks) 24 hours after injury induction (B). Nuclei of larvae treated with NFP + NM_187 appear healthy again, and primary processes are visible (C). C: Capillary; P: Podocyte. Scale bars in whole glomeruli images: 10 µm, Scale bars in inserts: 5 µm.

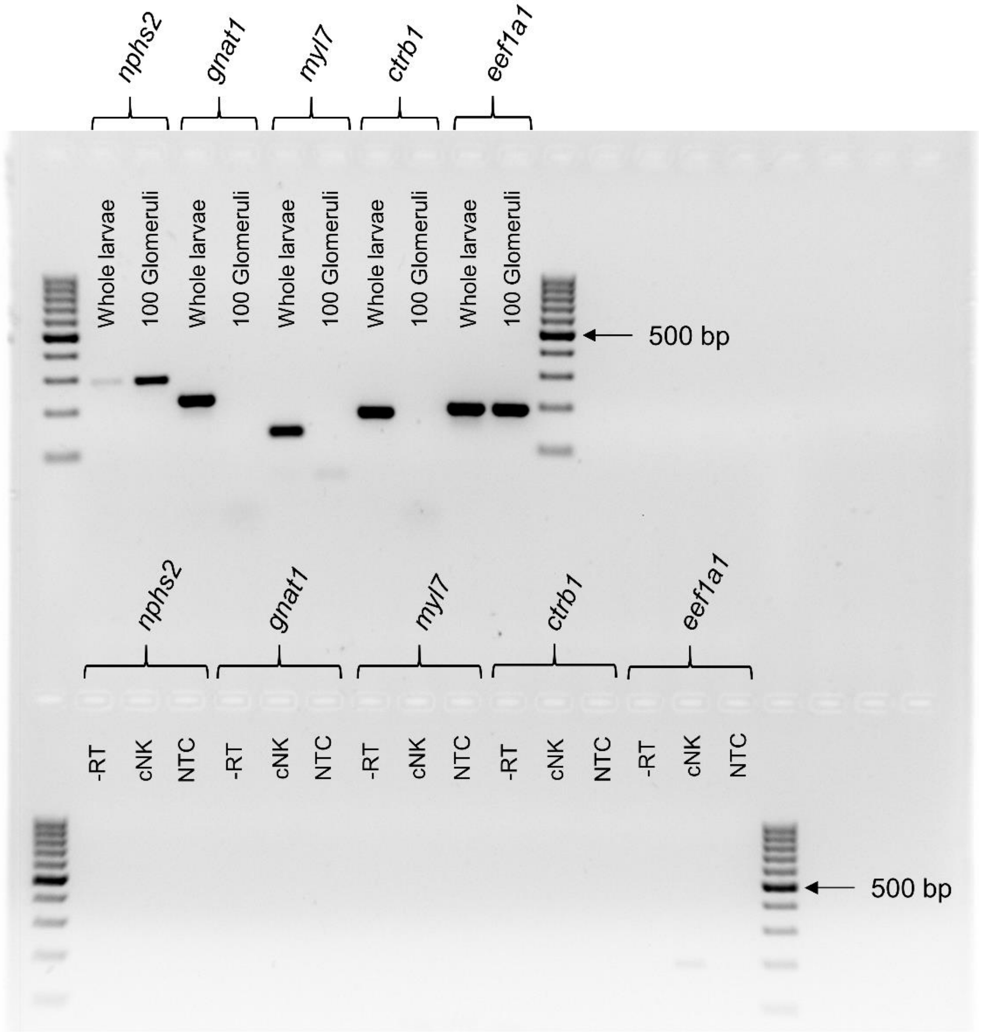


**Figure S5: Corresponding uncropped RT–PCR gel image.** The image shows the full agarose gel (2%) for the RT-PCR presented in Figure 5. The top row indicates the primer sets used and compares input from ten whole larvae and 100 isolated glomeruli. For all primer pairs, three controls were included: -RT (no reverse transcriptase during cDNA synthesis), cNK (no RNA during cDNA synthesis), and NTC (no-template control; water as PCR template).

### Supplemental Tables

**Table 1: Primers used for RT-PCR and RT-qPCR.**

| Gene | Tissue | Direction | Sequence | Amplicon |
|---|---|---|---|---|
| *eef1a1l1* | Housekeeping | Forward 5′–3′ | AAGGAGGGTAATGCTAGCGG | 190 kb |
| *eef1a1l1* | Housekeeping | Reverse 3′–5′ | GGGCGAAGGTCACAACCATA | 190 kb |
| *nphs2* | Podocytes | Forward 5′–3′ | GGCCCTGGGCTGATGTTTTA | 283 kb |
| *nphs2* | Podocytes | Reverse 3′–5′ | GAGCAATGCGTTTCCTGTCC | 283 kb |
| *mCherry* | Podocytes | Forward 5′–3′ | GAAGAAGACCATGGGCTGGG | 123 kb |
| *mCherry* | Podocytes | Reverse 3′–5′ | TTGACCTCAGCGTCGTAGTG | 123 kb |
| *gnat1* | Eye | Forward 5′–3′ | TCGCTGCTCTGAGTGCATAC | 218 kb |
| *gnat1* | Eye | Reverse 3′–5′ | GAAGGTGTTGGGACCGTCAT | 218 kb |
| *myl7* | Heart | Forward 5′–3′ | CAGACCCAGAGGAAACCATCC | 132 kb |
| *myl7* | Heart | Reverse 3′–5′ | GGTCAACCTCTTCTGCTGTGA | 132 kb |
| *ctrb1* | Pancreas | Forward 5′–3′ | GGCTATTCAGACCATTGCCG | 179 kb |
| *ctrb1* | Pancreas | Reverse 3′–5′ | TCACACACTTCATGCCACCA | 179 kb |

### Supplemental Movie Captions

Movie S1:

**Optical sections through a larval zebrafish glomerulus.** Podocytes are endogenously labeled with eGFP (green), and the slit diaphragm is detected with a nephrin antibody (magenta). Nuclei are shown in blue. The stack consists of 30 slices with 1 μm spacing and is recorded from ventral to dorsal at 2 frames per second. The larval glomerulus receives blood supply from the dorsal aorta, and the trench of this large vessel is visible in the last dorsal frames.

Movie S2:

**Reconstruction of larval podocytes.** This movie shows a rotation of all podocytes from a single glomerulus in a zebrafish larva. Podocytes are shown in green (eGFP) and major processes are visible.

Movie S3:

**Reconstruction of the slit diaphragm.** GLOMAGE enables visualization of the entire slit diaphragm in glomeruli in 3D. The slit diaphragm protein nephrin (magenta) bridges the filtration slits between podocyte foot processes.

Movie S4:

**Visualization of the proximal convoluted tubule.** Co-isolated tubules attached to the glomerulus enable 3D visualization of the tubule structure and labeling of the brush border. Podocytes are shown in green (eGFP), F-actin is detected with 543-conjugated phalloidin (magenta), and nuclei are shown in blue. Cells of the neck segment, which connect the glomerulus to the tubules, display fewer apical microvilli compared to proximal tubule epithelial cells.

Movie S5:

**Optical sections through a young (6 months) mouse glomerulus.** A nephrin antibody labels the slit diaphragm (magenta), a Dach1 antibody detects all podocyte nuclei (yellow) and the nuclear countersatin was performed with DAPI (blue). The stack is composed of 17 slices with 1 μm spacing and runs at 2 frames per second.

Movie S6:

**Three-dimensional reconstruction of a young mouse glomerulus.** The reconstruction shows the three-dimensionality of the glomerulus, nephrin is depicted in yellow, Dach1 in magenta and nuclei in blue.
